# Defining a functional hierarchy of millisecond time: from visual stimulus processing to duration perception

**DOI:** 10.1101/2025.06.06.658257

**Authors:** Valeria Centanino, Gianfranco Fortunato, Domenica Bueti

**Affiliations:** International School for Advanced Studies (SISSA), Trieste, Italy

## Abstract

The neural processing of subsecond durations recruits a wide network of areas. Although unimodal tuning has been shown in many of these regions, its role and link to perception remain unclear. Here, we used 7T functional MRI while participants performed a visual duration discrimination task to characterize unimodal responses along the cortical hierarchy. We found topographically-organized neuronal populations tuned to all presented durations in parietal and premotor cortices, and in the caudal supplementary motor area (SMA). In contrast, rostral SMA, inferior frontal cortex, and anterior insula showed neuronal preferences centered around the mean duration, which correlated with the boundary duration participants employed in the task. These differences suggest specialized roles of duration tuning across cortical regions —from discrete to categorical and subjective duration representations. Finally, correlations of neuronal preferences across areas highlighted a hierarchical organization of duration tuning. Together, our findings provide a mechanistic framework for duration perception in vision.

## Introduction

Time is a pervasive dimension of our everyday experience, and our continuous interaction with the environment relies on our ability to track and anticipate events unfolding within hundreds of milliseconds. In the past decades, a significant body of research in humans has shown that the processing of millisecond time engages several cortical and subcortical brain areas. Most of these brain regions seem engaged in temporal computations across a variety of tasks, sensory modalities and temporal ranges[1]–[3]. Beside the identification of this “timing network”, animal electrophysiology[4]–[10] and human functional magnetic resonance imaging (fMRI)[11]–[15] studies indicate that the brain processes millisecond time through neuronal units tuned to specific durations (i.e., exhibiting a unimodal response function). In humans, these neuronal units, which have been identified across a wide cortical network spanning lateral occipital, parietal, and frontal areas, are topographically organized in maps (i.e., neuronal units with similar duration preferences are spatially contiguous on the cortical surface)[11]–[15]. Interestingly, there is some overlap between the areas where temporal maps (“chronomaps”) have been observed and those previously identified as part of the timing network. However, the functional relationship between duration-tuned neural populations and timing areas remains elusive. In the first place, not all the areas of the timing network show unimodal tuning. For example, unimodal tuning has never been documented in the anterior insula, which is consistently activated by timing tasks[1]–[3] and considered central to the embodied account of time[2], [3], [16]–[18]. Second, unimodal tuning is not consistently reported in some brain areas. For instance, in the supplementary motor area (SMA), an area widely recognized for its central role in temporal computations[1], [3], [19] and where unimodal responses to stimulus durations have been recorded in monkeys[9], [10], [20], chronomaps are not always observed in humans[11]–[13]. Third, the redundancy of chronomaps across brain areas and their functional contribution to time processing and perception remains unclear. Evidence supporting a structured cortical organization of duration tuning, that also reflects functional differentiation, comes from two recent fMRI studies that focused on the visuo-spatial hierarchy[14], [15]. These studies showed that unimodal responses to duration gradually emerges from extrastriate visual areas onward, while at the earliest stages of visual processing, responses to durations are monotonic. This cortical organization, characterized by a shift from monotonic to unimodal duration tuning, suggests a transition in temporal processing: from an initial stage rooted in visual coding, where temporal information is extracted from the incoming event, to a subsequent stage dedicated to reading out that information. However, it remains unclear whether further changes in duration tuning occur along the cortical hierarchy to support other stages of temporal processing and, ultimately, shape duration perception. An indirect measure of the link between duration tuning and perception was reported by Hayashi and Ivry, who combined a duration adaptation paradigm with fMRI[21]. The authors found that, in the right supramarginal gyrus, durations similar to the adaptor stimulus evoked an attenuated response, consistent with the presence of duration-tuned populations that become suppressed following adaptation. Importantly, this modulation of brain response predicted the behavioral bias, supporting a connection between duration-tuned responses and perception. Further evidence for this link comes from studies showing a task dependence of duration-tuned populations in SMA. Duration maps in this area partially reshape when the tested duration range changes[11] or are not observed when participants were not actively engaged in a task[12], [13]. This latter finding is consistent with observations in other types of neural maps, whose presence depends on task demands. For instance, retinotopic maps in frontoparietal cortices are systematically and robustly elicited only when visual field mapping is combined with cognitively demanding tasks engaging attention or memory[22]–[24]. Overall, while existing evidence suggests a degree of dependence between duration-tuned populations and timing behavior, their direct relationship remains to be explored.

In summary, it remains unclear whether unimodal duration tuning changes across timing areas in a way that serves distinct functional roles, follows specific dependencies across areas, or directly contributes to perception. As a result, evidence has yet to converge into a comprehensive framework that accounts for the full transformation of millisecond durations from a stimulus feature (i.e., input) into a flexible perceptual object for decision-making and behavior (i.e., output).

To address these issues, in this study we asked 13 healthy human participants to perform a visual duration discrimination task while we measured brain activity using ultra-high-field (7T) fMRI. We then combined the classical fMRI localization approach with neuronal-based modeling to identify and characterize duration-tuned populations across a multitude of timing areas defined with high anatomical precision. Specifically, we aimed to determine (i) how the properties of unimodal tuning change across the cortical hierarchy; (ii) whether these changes are related across different areas, and (iii) whether they are linked to duration perception. Ultimately, our goal was to define a functional cortical hierarchy for temporal processing and perception in the visual modality.

## Results

To conduct this study, we used fMRI data already available within our research group[15], which met our requirements in terms of experimental manipulation and MRI acquisition (i.e., spatial and temporal resolution, brain coverage). In the experiment, participants were presented with a visual stimulus (i.e., a circular colored Gaussian noise patch subtending 1.5^◦^ of visual angle) whose display duration varied pseudo-randomly between 0.2 and 0.8 s. Participants were asked to judge whether the presented duration (i.e., the comparison duration) was longer or shorter than a previously internalized reference duration of 0.5 s. Stimuli were displayed in the lower half of the visual field, either 0.9^◦^ or 2.5^◦^ of visual angle to the left or right of a central fixation cross, and their spatial location was task irrelevant. Figure 1a shows a schematic representation of the trial structure with a highlight of the 4 spatial positions. Each participant completed 10 blocks (48 trials per block, 2 trials for each combination of stimulus duration and position) acquired in separate fMRI runs.

**Figure 1:**
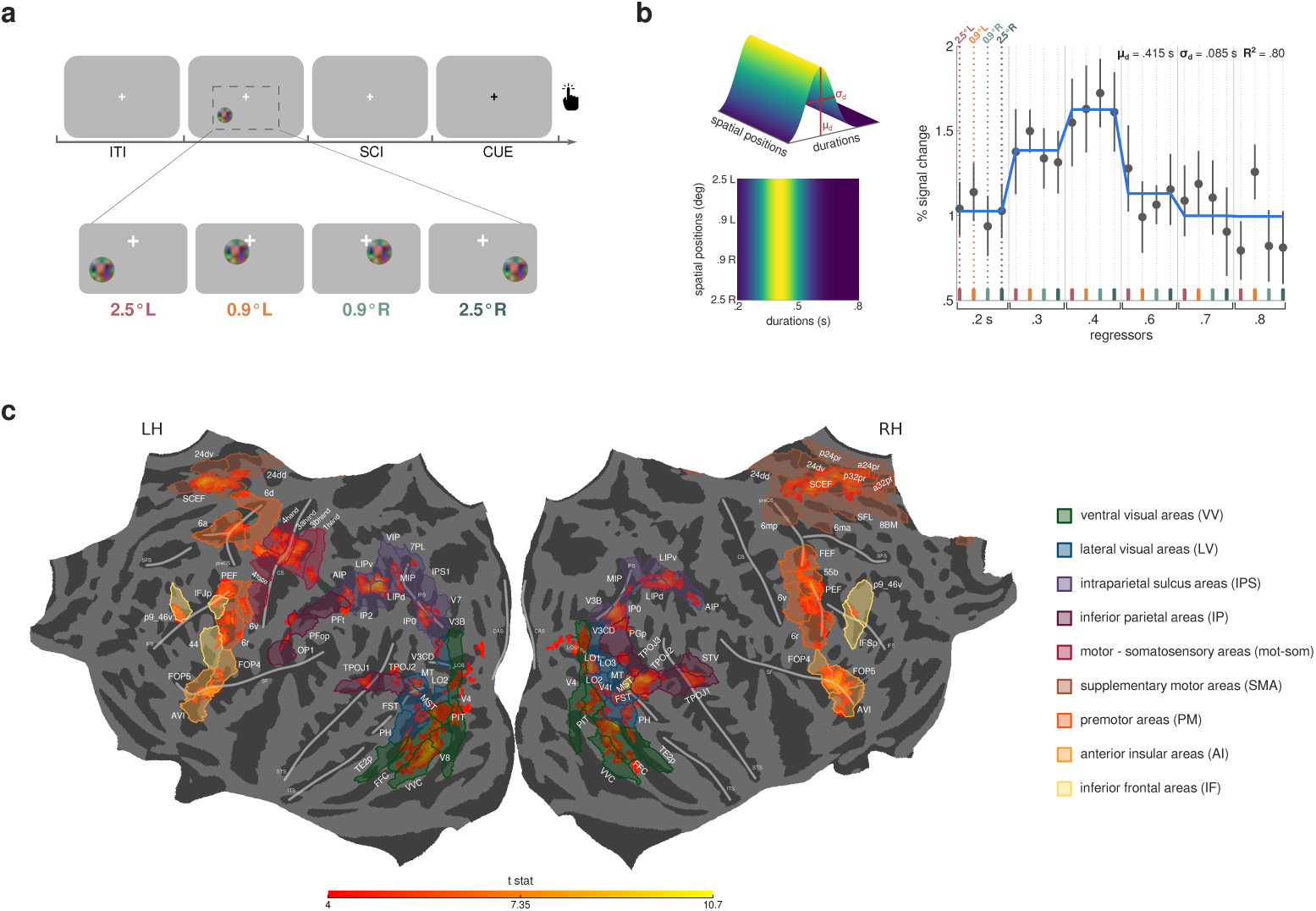
Experimental procedure, pRF modeling, and regions of interest (ROIs). **(a)** Schematic representation of the trial structure. In each trial, one of six different comparison durations (i.e., 0.2, 0.3, 0.4, 0.6, 0.7, 0.8 s) was presented at a specific location on the screen, which could be either 2.5*^◦^* or 0.9*^◦^* of visual angle in the lower-left (L) or lower-right (R) quadrant of the visual field, as shown in the close-up. Durations varied pseudo-randomly across trials, whereas positions varied sequentially, from 2.5*^◦^* L to 2.5*^◦^* R and back. Participants compared the comparison duration to a reference duration of 0.5, which they internalized during the training, and reported by a button press whether the comparison was longer or shorter than the reference. After a randomized interval from the offset of the comparison stimulus (stimulus-cue interval - SCI, uniformly drawn between 0.9-1.2 s), the response was cued with a color switch in the fixation cross from white to black. Trials were interleaved by a uniformly distributed inter-trial interval (ITI) spanning from 1.8 to 2.5 s. The fixation cross was displayed at the center of the screen throughout the experiment. See *Methods - Stimuli and procedure*. **(b)** Result of the pRF modeling in one representative vertex. On the left, three-dimensional (top) and two-dimensional (bottom) representations of the pRF model are shown. Model parameters, i.e., its preference (*µ_d_*) and its sensitivity (*σ_d_*), are highlighted in the three-dimensional representation. On the right, the plot shows the model fit. The solid line indicates the model prediction. Black dots represent the median GLM beta weights in percentage of signal change for each combination of stimulus duration and spatial position (color-coded), with error bars indicating standard errors. The top right corner of the plot reports the parameters and goodness of fit of the model prediction. See *Methods – Population receptive field (pRF) modeling*. **(c)** Regions of interest (ROIs) are displayed on a common (fsaverage) flattened cortical surface, overlaid on group-level t-value clusters (p*<*0.001 FWE-corrected for multiple comparisons at the cluster level) obtained from a GLM analysis that identified cortical locations significantly activated at the offset of the 6 presented durations. T-statistics values are color-coded from red (t-stats = 4) to yellow (t-stats = 10.7). The HCP MMP 1.0 atlas[25] and the topological atlas by Sereno and colleagues[26] were used to localize the t-value clusters, resulting in 47 ROIs in the left hemisphere and 46 ROIs in the right, with 29 ROIs shared between hemispheres. See *Methods - Regions of interest (ROIs) identification*. ROIs are labeled in white and follow the nomenclature of their respective atlases. They are color-coded according to functional streams: green for ventral visual (VV), blue for lateral visual (LV), violet for IPS, purple for inferior parietal (IP), red for motor-somatosensory (mot-som), brown for SMA, orange for premotor (PM), ochre for anterior insula (AI), yellow for inferior frontal (IF). Major sulci are displayed as thick semi-transparent white lines, with the following abbreviations: CAS = calcarine sulcus, LOS = lateral occipital sulcus, ITS = inferior temporal sulcus, STS = superior temporal sulcus, IPS = intraparietal sulcus, SF = Sylvian fissure, CS = central sulcus, IFS = inferior frontal sulcus, SFS = superior frontal sulcus, preCS = precentral sulcus.

We conducted a first-level general linear model (GLM) analysis (see *Methods – General linear model (GLM) analysis*), modeling the offsets of the 24 unique combinations of comparison durations and positions as events of interest. The resulting GLM beta weights were then used to feed a vertexwise modeling procedure based on the population receptive field (pRF) approach[27]. To specifically target neuronal populations selective for temporal information regardless of spatial information, we designed a model that assumes unimodal tuning for stimulus duration, and is invariant to stimulus position (see *Methods – Population receptive field (pRF) modeling*). The model describes the BOLD response of each vertex with two parameters: *µ_d_*, the duration evoking the greatest neuronal response (i.e., duration preference), and *σ_d_*, the sensitivity of the response. Figure 1b illustrates an example of the model’s response function, alongside the corresponding pRF modeling result. All the analyses presented in this work focused on the duration preference (*µ_d_*) parameter.

We restricted our analyses to a set of cortical regions of interest (ROIs) that, according to a group-level GLM analysis, showed a significant activation at the offset of the 6 presented durations (activations were p*<*0.001 FWE-corrected for multiple comparisons at cluster-level). Locations were identified employing two highly-parcellated atlases (see *Methods – Regions of interest (ROIs) identification*). This approach enabled us to explore all cortical locations recruited for duration processing in our experimental paradigm with high anatomical precision. To provide a comprehensive picture, we also grouped the ROIs into 9 functional streams and conducted all subsequent analyses at both stream and ROI levels. We identified 6 areas in the ventral visual (VV) stream (bilateral V4, left V8, bilateral PIT, bilateral FFC, bilateral VVC, bilateral TE2p); 9 areas in the lateral visual (LV) stream (bilateral V3CD, right LO1, bilateral LO2, right LO3, right V4t, bilateral MT, bilateral MST, bilateral FST, bilateral PH); 11 areas within or adjacent to the IPS (bilateral V3B, left V7, bilateral IP0, left IPS1, left 7PL, left VIP, bilateral MIP, bilateral LIPv, bilateral LIPd, left IP2, bilateral AIP); 8 areas in the inferior parietal (IP) lobule (right PGp, right TPOJ3, bilateral TPOJ2, bilateral TPOJ1, right STV, left PFt, left PFop, left OP1); 5 motor and somatosensory (mot-som) areas, all in the left hemisphere (area 1 - hand subdivision, area 3b - hand subdivision, area 3a - hand subdivision, area 4 - hand subdivision, area 4 - face subdivision); 11 areas in the SMA (right 6mp, right 6ma, bilateral 24dd, bilateral 24dv, right p24pr, right SFL, bilateral SCEF, right 8BM, right p32pr, right a32pr, right a24pr); 7 premotor (PM) areas (left 6d, left 6a, right FEF, right 55b, bilateral PEF, bilateral 6v, bilateral 6r); 3 areas in the anterior insular (AI) region (bilateral FOP4, bilateral FOP5, bilateral AVI); 4 areas of the inferior frontal (IF) lobule (left IFJp, left 44, right IFSp, bilateral p9 46v). Figure 1c shows all the ROIs on a common surface, color-coded according to the functional stream they belong to.

### How duration preferences change along the cortical hierarchy

To obtain an initial indication of changes in duration tuning along the cortical hierarchy, we assessed how preferred durations vary across functional streams and ROIs (see *Methods - Analysis of duration preference changes along the cortical hierarchy*). Figure 2 shows preferred duration maps for a selection of ROIs across all participants, while figure 3 illustrates the vertex-wise distributions at group-level of preferred durations for each functional stream (a) and ROI (b).

**Figure 2:**
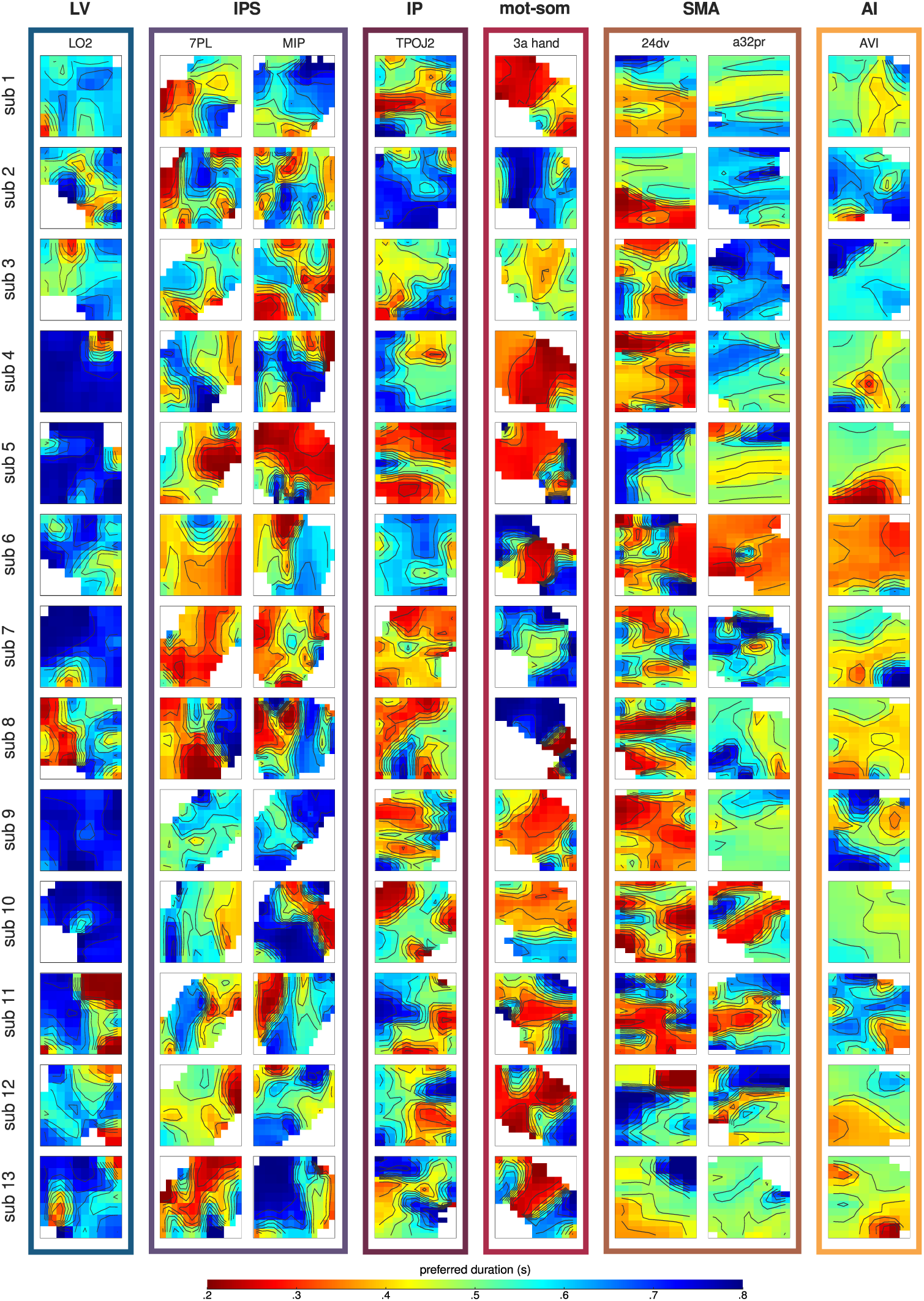
Maps of duration preferences. Duration preference maps are shown for all participants (in separate rows) across a selection of ROIs (ordered from occipital to frontal in the different columns), in either the left (LO2, 7PL, MIP, TPOJ2, 3a_hand, AVI) or the right (24dv, a32pr) hemisphere. Maps belonging to the same functional stream are grouped with a color-coded outline (blue for LV, violet for IPS, purple for IP, red for mot-som, brown for SMA, ochre for AI). Duration preferences are color-coded from red (.2 s) to blue (.8 s). Black lines indicate isolines that join locations with equal preference values.

**Figure 3:**
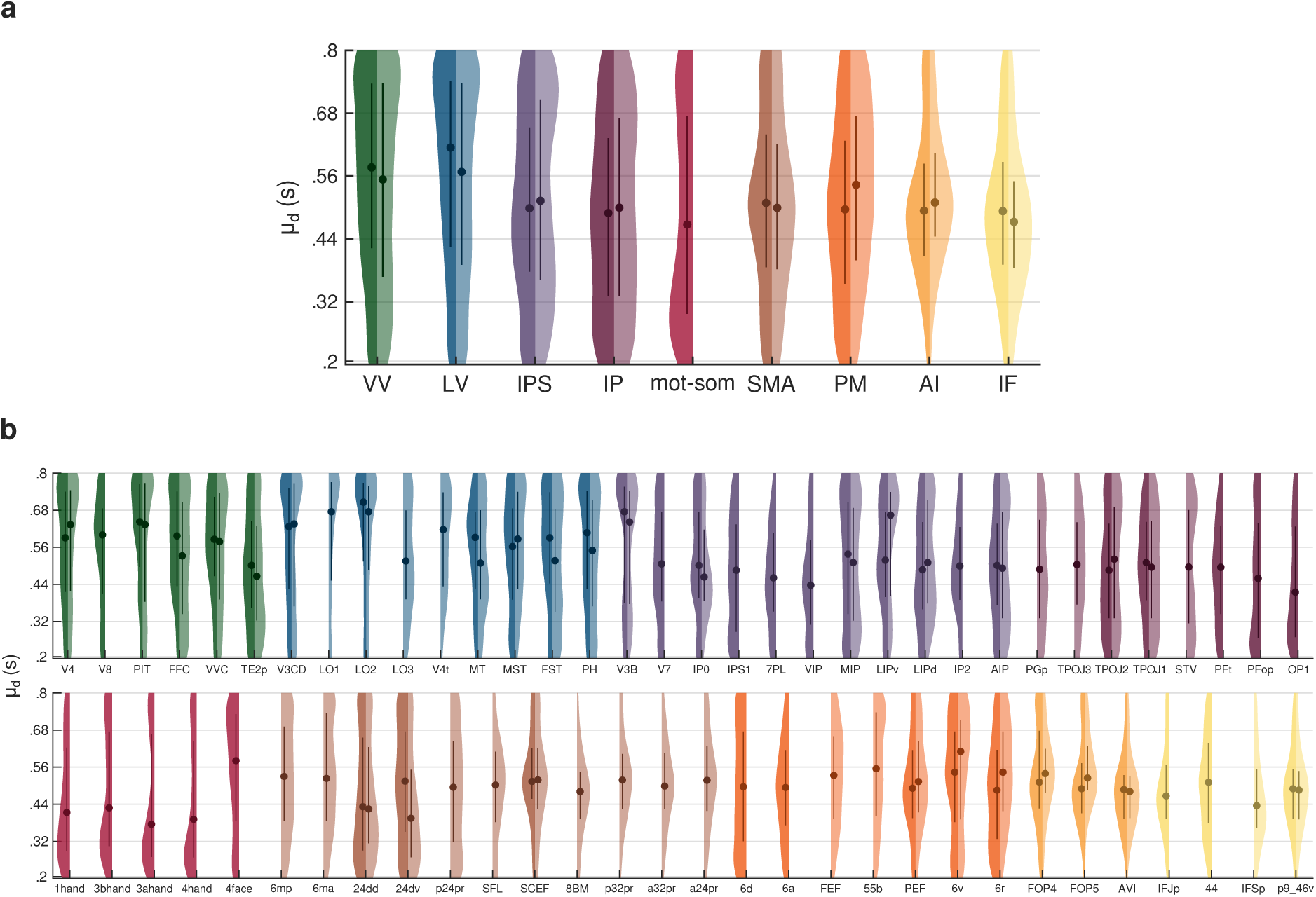
Group-level distributions of duration preferences. Each violin plot shows the vertex-wise distribution of duration preferences at group-level across streams **(a)** and ROIs **(b)**. Streams and ROIs are ordered from occipital to frontal and from dorsal to ventral. Violins are color-coded according to functional streams: green for VV, blue for LV, violet for IPS, purple for IP, red for mot-som, brown for SMA, orange for PM, ochre for AI, yellow for IF. The left side of the violins (darker shades) represents the left hemisphere, while the right side (lighter shades) represents the right hemisphere. Dots indicate the median of each distribution, thick lines represent the interquartile range. The kernel density estimate of each distribution was computed using a 10% bandwidth. See *Methods - Analysis of duration preference changes along the cortical hierarchy*.

To quantify changes in duration preferences across functional streams, we tested a linear mixed effect (LME) model with stream as factor, and ROI and participant as random intercepts, using individual median *µ_d_* for each ROI (model formula: *µ_d_* ∼ *stream* + (1|*ROI*) + (1|*subjectID*), marginal *R*^2^: 0.09, conditional *R*^2^: 0.16). Type III ANOVA on model estimates revealed a main effect of stream (F(8,55) = 6.73 p *<* 0.0001). Specifically, the median preferred duration was higher in occipital visual streams compared to parietal and frontal ones (all t(55) *>* 4.62, p *<* 0.001 comparing LV to IPS, IP, mot-som, SMA, and IF; all t(55) *>* 3.44, p *<* 0.05 comparing VV to mot-som and IF). Similarly, to quantify changes in duration preferences across ROIs, we tested a LME model with ROI as factor and participant as random intercept, using individual median *µ_d_* for each ROI (model formula: *µ_d_* ∼ *ROI* + (1|*subjectID*), marginal *R*^2^: 0.18, conditional *R*^2^: 0.20). Type III ANOVA on model estimates revealed a main effect of ROI (F(63,756) = 2.98 p *<* 0.0001). In LO1 and LO2 (belonging to the LV stream) the median *µ_d_* was significantly higher than in several ROIs within parietal, supplementary motor, and inferior frontal regions (all t(756) >4.55, p *<* 0.05 comparing LO1 to 7PL, VIP, OP1; all t(756) *>* 4.32, p *<* 0.05 comparing LO2 to IPS1, 7PL, VIP, OP1, 8BM, 24dd, 24dv, IFJp, IFSp, p9 46v). In motor and somatosensory hand areas, the median *µ_d_* was significantly lower than in many occipital visual areas (all t(756) *>* 4.31, p *<* 0.05 comparing areas 1 hand, 3a hand, and 4 hand to V4, PIT, V3CD, LO1, LO2, V4t, and V3B). The full set of statistics is reported in supplementary tables 1 - 4. All reported p-values were Bonferroni-corrected for multiple comparisons.

In line with the statistical assessments, the *µ_d_* distributions shown in figure 3 are mainly skewed towards longer preferences in the ventral and lateral visual streams (VV and LV), whereas they shift towards shorter preferences in motor and somatosensory cortices (mot-som). Interestingly, in the IPS and inferior parietal (IP) areas, the distributions are more evenly spread across the duration range, while from SMA onward they are mainly centered around the mean of the duration range. At the ROI level, this pattern remains evident (see also figure 2), although not all ROIs within each stream show consistency. The highest variability among ROIs was observed in the IPS, IP, and SMA streams. These findings highlight significant changes in the distribution of duration preferences across areas, suggesting that response properties of duration-tuned populations may vary throughout the cortical hierarchy.

### Categorization of duration preferences along the cortical hierarchy

The previous analysis of *µ_d_* distributions suggested that a key difference between cortical areas may be the temporal range they are tuned to (i.e., either the full range of presented durations or a narrower range around a specific duration). These different ranges of duration selectivity may thus indicate distinct processing stages of stimulus temporal information. To address this, we grouped duration preferences into 5 categories, each corresponding to a different portion of the tested range: short (.2-.32 s), mid-short (.32-.44 s), medium (.44-.56 s), mid-long (.56-.68 s), and long (.68-.8 s) (see *Methods - Analysis of duration preference categories along the cortical hierarchy - Categorization of duration preferences*). For each participant, we then computed the fraction of vertices within each duration category for each functional stream and ROI, averaging across hemispheres for bilateral ones. Figure 4 shows the group-level fractions of vertices across streams (a) and ROIs (c). These data were analyzed using two separate two-way repeated-measures ANOVA, with category and either stream or ROI as within-subject factors. The ANOVA showed a significant main effect of category (across streams: F(4,48) = 5.45, p *<* 0.005; across ROIs: F(4,48) = 4.38, p *<* 0.005) and a significant interaction between category and either stream or ROI (across streams: F(32,384) = 6.40 p *<* 0.0001; across ROIs: F(252,3024) = 3.15 p *<* 0.0001), indicating that duration categories are represented differently across brain regions. As also shown in figure 4a, in the ventral and lateral visual streams (VV and LV), the long category was the most represented among all others (t(540) *>* 3.39, p *<* 0.01 in VV comparing long with short, mid-short, and medium; all t(540) *>* 4.06, p *<* 0.001 in LV), while in the anterior insula (AI) and inferior frontal cortex (IF), the medium category prevailed (all t(540) *>* 4.60, p *<* 0.0001 in AI; t(540) >4.93, p *<* 0.0001 in IF comparing medium with short, mid-long, and long). In the motor and somatosensory areas (mot-som) instead, the short category prevailed over the medium and the mid-long ones (t(540) *>* 4.24, p *<* 0.0005). Interestingly, in the other streams, all duration categories were equally represented. A similar pattern of results was observed at the ROI level, although the finer resolution allowed us to identify some exceptions. In V3B -the most occipital ROI of the IPS stream, located at its posterior limit- the long category was the most represented, similar to the ROIs in the ventral and lateral visual streams (all t(3840) *>* 4.20, p *<* 0.0005). Likewise, in OP1, a region of the parietal operculum likely contributing to somatosensory processing[28], [29], short durations were prevalent, resembling motor and somatosensory areas (t(3840) *>* 2.94, p *<* .05 comparing short with medium and mid-long). Another interesting observation concerns the ROIs within SMA, which appeared to separate into two subgroups. Rostral ROIs mainly showed preferences for the medium range (SCEF: t(3840) *>* 3.02, p *<* 0.05 comparing medium with short, mid-short, and mid-long; SFL: t(3840) = 2.93, p *<* 0.05 comparing medium with long; 8BM: t(3840) *>* 3.62, p *<* 0.005 comparing medium with short, mid-long, and long; p32pr: t(3840) >3.17, p *<* 0.05 comparing medium with short, mid-short, and long; a32pr: t(3840) *>* 3.33, p *<* 0.01 comparing medium with short, mid-short, and long; a24pr: t(3840) = 3.80, p *<* 0.005 comparing medium with short), while caudal ROIs showed no significant differences across duration categories. The complete set of statistics is reported in supplementary tables 5 - 12. All reported p-values were Bonferroni-corrected for multiple comparisons. These findings indicate that, along the cortical hierarchy, different areas are selective to different duration ranges. In occipital visual areas, preferences for long durations likely reflect a monotonic tuning for stimulus duration, as already reported in previous studies[14], [15], [30], which may mediate the encoding of temporal information. In parietal, premotor, and caudal supplementary motor areas, preferences across the full range of presented durations suggest instead the presence of duration-tuned populations that read out the temporal information at hand[11], [15]. Finally, inferior frontal regions, anterior insula and the rostral subdivisions of SMA, by representing the mean of the duration range, may provide the categorical boundary employed for the discrimination task[31]. In motor and somatosensory areas, the selectivity for short durations is likely a by-product of motor preparation responses, as argued in the Discussion section. Collectively, these findings point to three distinct stages of temporal processing, each characterized by different properties of duration tuning and implemented in specific cortical areas: duration encoding, duration readout, and duration categorization.

**Figure 4:**
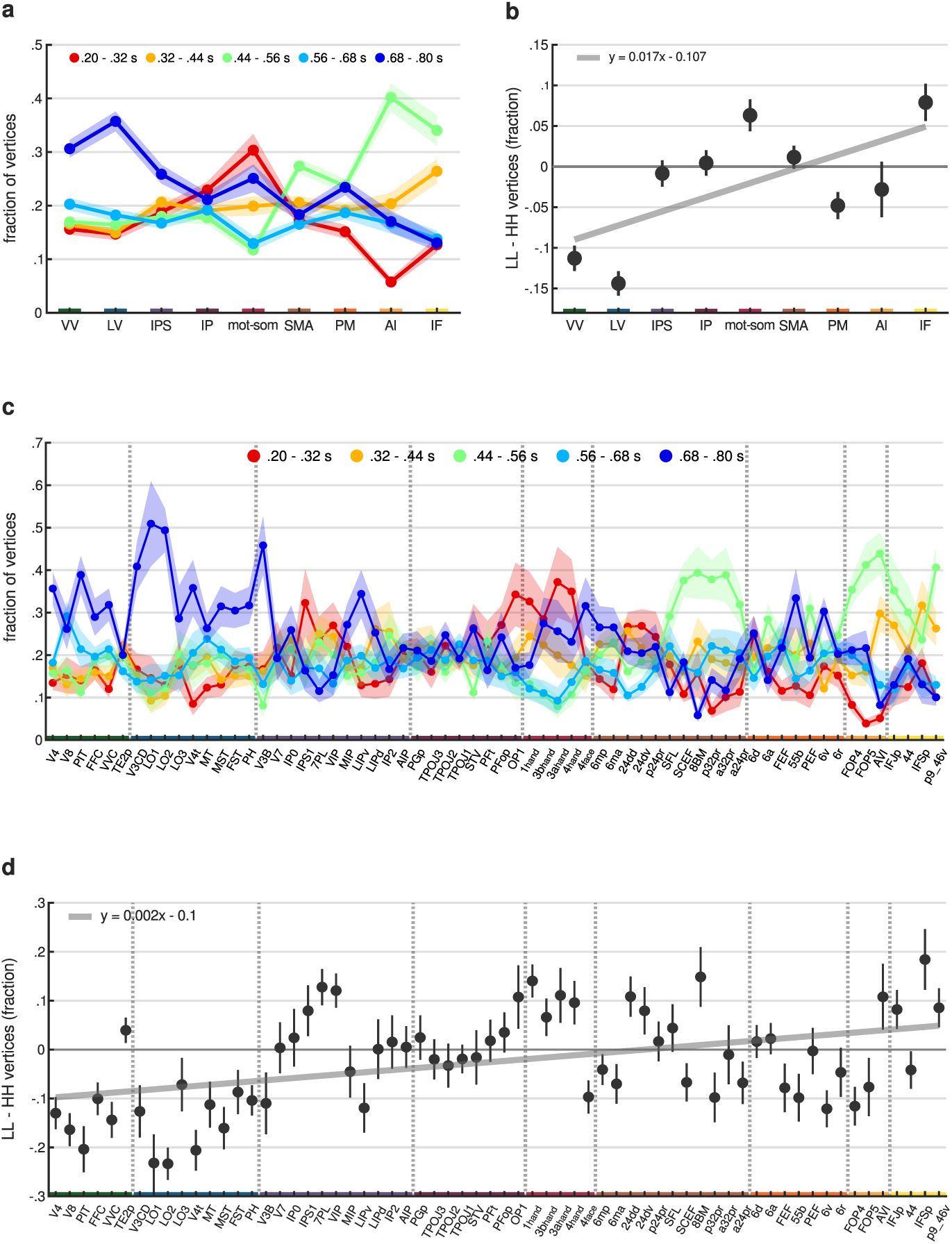
Categorization and local Moran’s I of duration preferences. Panels **(a)** and **(c)** show the group-level fractions of vertices (y-axis) assigned to each duration category across streams and ROIs (x-axis), respectively. Each solid line represents a different duration category: red for short (.2-.32 s), yellow for mid-short (.32-.44 s), green for medium (.44-.56 s), light-blue for mid-long (.56-.68 s), blue for long (.68-.8 s). Dots indicate the mean fraction across participants and hemispheres, with the shaded area representing standard errors. Panels **(b)** and **(d)** show the group-level differences (y-axis) in the faction of vertices between low-low (LL) and high-high (HH) spatial associations across streams and ROIs (x-axis), respectively. These spatial associations were computed using the local Moran’s I statistic. Dots represent the mean difference across participants and hemispheres, with error bar indicating standard errors. Regression lines are displayed in gray, and their equations are shown in the top left corner of the corresponding plot. Across all panels, streams and ROIs are ordered from occipital to frontal and from dorsal to ventral. Streams are color-coded on the x-axis: green for VV, blue for LV, violet for IPS, purple for IP, red for mot-som, brown for SMA, orange for PM, ochre for AI, yellow for IF. Vertical dashed lines in panels c and d separate different streams. See *Methods - Analysis of duration preference categories along the cortical hierarchy*.

To further validate the previous analysis, we explored the spatial association between neighboring duration preferences by computing the local Moran’s I statistic. This statistic is computed, for each vertex in the brain, as the correlation between its duration preference and the average preference of its neighbors. In this way, it identifies vertices whose duration preference is significantly associated with their neighbors and classifies the resulting association into the following types: high duration preference surrounded by high ones (high-high, or HH); low surrounded by low (low-low, or LL); high surrounded by low (high-low, or HL); and low surrounded by high (low-high, or LH). Before computing the Moran’s I, we centered duration preference values on the mean preference of the hemisphere. Therefore, the terms “low” and “high” are relative to this average, regardless of the absolute deviation. For further details, see *Methods - Analysis of duration preference categories along the cortical hierarchy - Local Moran’s I*. We hypothesized that brain regions selective to one specific duration category (i.e., VV, LV, mot-som, rostral SMA, AI, IF) would also show one dominant type of spatial association between neighboring vertices. On the other hand, regions where all duration categories are equally represented (i.e., IPS, IP, caudal SMA) should show multiple spatial association types. To test this, for each participant we computed the fraction of vertices showing each association type within each ROI. Interestingly, the HL and LH associations were nearly absent. On average across participants, hemispheres and ROIs, their fractions were less than 0.019 (see also supplementary figure 1), indicating that similar duration preferences tend to cluster together. For this reason, we focused the following analyses on the LL and HH associations only, comparing their respective fraction of vertices. Figure 4 shows the group-level differences across streams (b) and ROIs (d). To assess changes in the LL-HH difference across functional streams, we used a LME model with stream as factor, and ROI and participant as random intercepts (model formula: *LL* − *HH* ∼ *stream* + (1|*ROI*) + (1|*subjectID*), marginal *R*^2^: 0.09, conditional *R*^2^: 0.15). Similarly, to quantify changes across ROIs, we used a LME model with ROI as factor and participant as random intercept (model formula: *LL* − *HH* ∼ *ROI* + (1|*subjectID*), marginal *R*^2^: 0.17, conditional *R*^2^: 0.19). Type III ANOVA on model estimates revealed a main effect of both stream (F(8,55) = 6.47, p *<* 0.0001) and ROI (F(63,756) = 2.81, p *<* 0.0001), indicating a systematic change in the LL-HH difference along the cortical hierarchy (see supplementary tables 13 and 14). Specifically, this difference increased along the cortical hierarchy, shifting from negative to positive values (*β* = 0.017, t(7)= 2.25, p = 0.059 across streams; *β* = 0.002, t(62) = 3.79, p *<* 0.0005 across ROIs). In occipital regions (nearly all ROIs within VV and LV), HH associations were more frequent than LL, whereas in inferior frontal and anterior insular regions (AVI, IFJp, IFSp, p9 46v) the opposite happened. ROIs in the motor-somatosensory stream also showed a prevalence of LL associations. Interestingly, several ROIs within the IPS, inferior parietal lobule (IP), and SMA showed a LL-HH difference close to 0, indicating an equal presence of both spatial associations. See supplementary tables 15 - 20 for additional supporting statistics. These results are in line with the previous findings from the categorization analysis of duration preferences. Areas selective for a specific duration category also show one main type of spatial association between neighboring vertices: HH in occipital visual areas, where long preferences predominated, and LL in motor-somatosensory regions and inferior frontal cortex, where preferences were mainly for low and medium durations, respectively. In regions showing the full range of duration preferences, such as parietal and supplementary motor regions, HH and LL associations coexisted. Overall, these results suggest that visual duration processing is supported by both duration selectivity and the local cortical arrangement of duration information. Importantly, different duration tuning properties emerge at different levels of the cortical hierarchy. These findings strengthens the idea that duration tuning contributes to the transformation and representation of duration information across the cortical hierarchy by mediating different stages of temporal processing.

### The topographic organization of duration preferences along the cortical hierarchy

A key property of unimodal responses to duration is their organization into topographic maps along the cortical surface[11]–[13]. Our next goal was to investigate whether and how the topographic properties of duration preferences vary across areas, gaining further details about the changes of unimodal tuning along the cortical hierarchy.

First, we tested whether duration preferences were spatially clustered within each ROI by computing the global Moran’s I statistic (see *Methods - Analysis of the topographic organization of duration preferences along the cortical hierarchy - Global Moran’s I*). The global Moran’s I quantifies the overall spatial autocorrelation of duration preferences within the ROI and is computed by averaging all local Moran’s I statistics (i.e., the correlation between each vertex’s duration preference and the average preference of its neighbors). A global Moran’s I close to 0 indicates a random distribution of duration preferences along the cortical surface. Values above or below zero indicate different degrees of spatial clustering, characterized by either a positive (i.e., clustering of similar duration preferences) or a negative (i.e., clustering of opposite preferences) spatial autocorrelation. Figure 5a shows the group-level distributions of this statistic across streams, while supplementary figure 2 shows group-level Moran’s I distributions across both streams (a) and ROIs (b). Due to the unique neighborhood configuration of each ROI, statistical comparison of Moran’s I values across ROIs is not feasible. The results revealed a positive and relatively stable Moran’s I across the cortical hierarchy (median across participants above 0.50 in all streams and above 0.34 in all ROIs), indicating that vertices with similar preferences are consistently clustered together. However, occipital visual areas (VV and LV) showed slightly lower values than the other regions.

**Figure 5:**
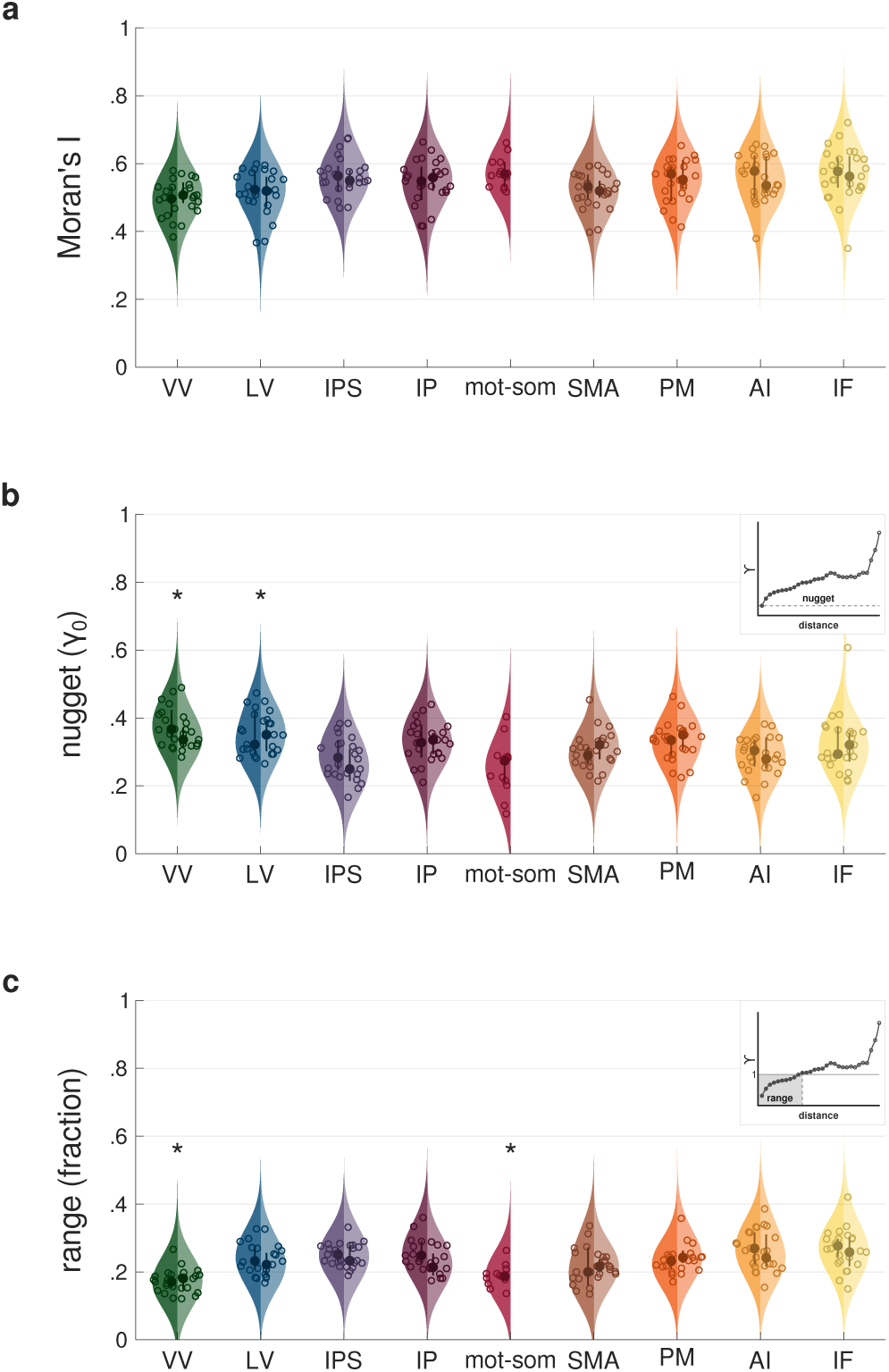
Group-level distributions of Moran’s I, nugget and range values. Each violin plot represents the group-level distribution of Moran’s I statistic **(a)**, variogram nugget **(b)** and range **(c)** values across streams. Nugget values are expressed as a fraction of the total variance within the ROI; range values are expressed as a fraction of the maximum distance between vertices of the ROI. Streams are ordered from occipital to frontal and from dorsal to ventral, and are color-coded as follows: green for VV, blue for LV, violet for IPS, purple for IP, red for mot-som, brown for SMA, orange for PM, ochre for AI, yellow for IF. The left side of each violin represents the left hemisphere (darker shades), while the right side represents the right hemisphere (lighter shades). Dots indicate the median of each distribution, while circles correspond to individual data points. Thick lines represent interquartile ranges. The kernel density estimates were computed using a 7% bandwidth. In panels b and c, asterisks indicate streams that differ statistically from the others. An example variogram highlighting the nugget and the range is shown as an inset in panels b and c. In the inset graph, each dot represents the variance (*γ*) in duration preferences between pairs of vertices at increasing distance. Dots shading reflects the number of vertex pairs at each distance, with darker shades indicating a higher count. The solid line (*γ* = 1) in c indicates the variance in duration preferences across all vertices, without accounting for spatial structure. The nugget (i.e., the variance at the shortest vertex distance) is marked in b by the dashed line, whereas in c the range (i.e., the distance at which the total variance is reached) is highlighted by a gray box. See *Methods - Analysis of the topographic organization of duration preferences along the cortical hierarchy*.

We next computed the experimental variogram for each ROI and extracted two additional indicators of spatial autocorrelation: the nugget and the range (see *Methods - Analysis of the topographic organization of duration preferences along the cortical hierarchy - Variogram*). The variogram estimates spatial autocorrelation by computing the variance among the duration preferences of vertices at varying distances within the ROI. In a variogram, the nugget is the variance observed at the shortest distance between vertices in the ROI (see the inset in figure 5b), and the range is the distance between vertices required to reach the overall variance of the ROI (see the inset in figure 5c). While the previously described global Moran’s I captures the overall spatial pattern within the ROI, the nugget provides finer resolution by focusing on neighboring vertices and reflects the strength of the spatial autocorrelation. The range, in contrast, quantifies the persistence of spatial autocorrelation along the cortical surface. Figure 5 shows the group-level distributions of nuggets (b) and ranges (c) across streams, while supplementary figures 3 and 4 show the group-level distributions of nuggets and ranges respectively, across both streams (b) and ROIs (c). To assess changes of nugget and range across functional streams, we tested two LME models with stream as factor and participant as random intercept. The LME model formulas were as follows: *nugget* ∼ *stream* + (1|*subjectID*) (marginal *R*^2^: 0.31, conditional *R*^2^: 0.57) and 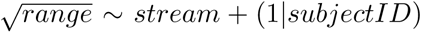 *stream* + (1 *subjectID*) (marginal *R*^2^: 0.51, conditional *R*^2^: 0.53). Type III ANOVA on model estimates revealed a main effect of stream for both nugget and range (nugget: F(8,96) = 10.26 p *<* 0.0001; range: F(8,96) = 15.76 p *<* 0.0001). Nuggets were significantly higher in occipital visual streams compared to parietal and frontal ones (all t(96) *>* 3.80, p *<* 0.01 comparing VV to IPS, mot-som, SMA, and AI; all t(96) >4.06, p *<* 0.01 comparing LV to IPS, mot-som, and AI). Conversely, ranges were significantly lower in both ventral visual (VV) and motor-somatosensory (mot-som) streams compared to the majority of the other streams (all t(96) *>* 4.14, p *<* 0.01 comparing VV to LV, IPS, IP, SMA, PM, AI, and IF; all t(96) *>* 4.14, p *<* 0.01 comparing mot-som to LV, IPS, IP, PM, AI, and IF). The complete set of statistics is reported in supplementary tables 21 - 24. All reported p-values were Bonferroni-corrected for multiple comparisons. Similar results were obtained performing LME model analyses at the ROI level (see supplementary tables 25 - 28).

In summary, the assessment of topographic properties of duration preferences with Moran’s I, nuggets, and ranges revealed that occipital (VV and LV) and motor-somatosensory areas exhibit a weaker spatial arrangement compared to the other areas. This suggests that in these regions the spatial organization of duration preferences is less critical for duration processing.

### Long-range relationships between duration preferences

After characterizing the changes in duration preferences across different brain regions, we next explored whether and how these changes are related along the cortical hierarchy. To address this at the ROI level, for each participant we computed the median duration preference within each ROI and then calculated the group-level Kendall correlation matrix, shown in figure 6 (see *Methods - Analysis of long-range relationships between duration preferences*). This analysis allowed us to capture the overall pattern of relationships between duration preferences across ROIs. From a qualitatively point of view, the correlation matrix showed mainly positive values within functional streams, with higher consistency in visual (VV, LV), motor-somatosensory (mot-som), and anterior insular (AI) streams. This indicates that duration preferences change similarly in ROIs that are in close anatomical and functional proximity. More interestingly, the correlation matrix allowed us to explore relationships between ROIs from distinct and distant streams, providing insight into the hierarchical organization of duration preference changes. We observed negative correlations between ROIs at the opposite ends of the hierarchy. Specifically, ROIs in the ventral visual stream (VV) were positively correlated with the nearby ROIs in the lateral visual stream (LV) but negatively correlated with all other ROIs from more distant streams. Similarly, ROIs rostral to the motor-somatosensory stream (mot-som) were positively correlated among themselves but negatively correlated with both occipital streams (VV and LV). For the parietal streams (IPS and IP), correlation patterns were more nuanced. ROIs in the IPS were positively correlated with occipital (VV and LV), anterior insular (AI) and inferior frontal (IF) regions, but mainly negatively correlated with the other streams. By contrast, ROIs in the inferior parietal cortex (IP) appeared to segregate into two groups. PGp and TPOJ3 showed overall weak correlations, but tended to positively correlate with some ROIs in the lateral visual (LV) stream. From TPOJ2, ROIs showed instead negative correlations with caudal ROIs and positive correlations with rostral ROIs.

**Figure 6:**
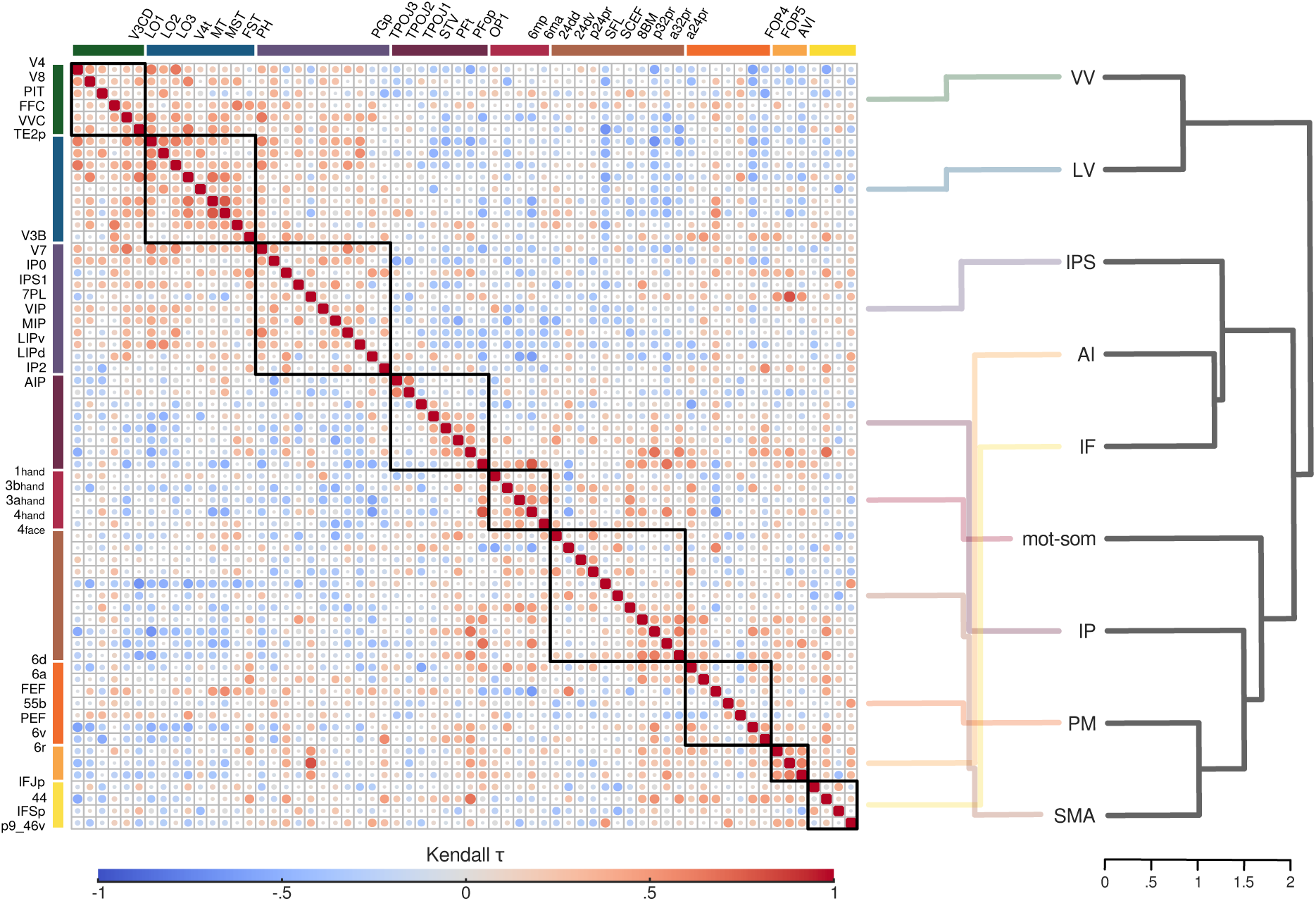
Long-range relationships between duration preferences. The left side shows the group-level Kendall correlation matrix, computed using the median preferred duration of each ROI and participant. The size and color of each dot represent the Kendall *τ* value. This visualization was generated using the corrplot package[32] in R. ROIs on x and y axes are arranged from occipital to frontal and from dorsal to ventral, with colors indicating their respective streams: green for VV, blue for LV, violet for IPS, purple for IP, red for mot-som, brown for SMA, orange for PM, ochre for AI, yellow for IF. Black squares within the matrix outline ROIs belonging to the same stream. The right side displays the dendrogram representation of the Kendall correlation matrix at the stream level. Semi-transparent lines, color-coded by stream, connect the ROI representation on the left to the stream representation on the right for visualization purposes. See *Methods - Analysis of long-range relationships between duration preferences*.

To better capture and quantify long-range relationships between duration preferences, we recomputed the Kendall correlation matrix using individual median preferred durations of each stream and used these data to build the dendrogram shown in figure 6 (see *Methods - Analysis of long-range relationships among duration preferences*). The dendrogram represents the similarity between streams based on the correlation of their duration preferences. Streams with stronger positive correlations are placed closer together, while those with weaker or negative correlations are located farther apart, resulting in a hierarchical representation of duration preference correlations. This analysis identified three main clusters. The first cluster included lateral and ventral visual streams (VV and LV). The second cluster included the IPS, the anterior insula (AI), and the inferior frontal cortex (IF). The third cluster comprised the inferior parietal lobule (IP), the medial and lateral premotor cortices (SMA and PM), and the motor-somatosensory cortices (mot-som). Within the second cluster, anterior insula and inferior frontal areas were more similar to each other than to the IPS. Within the third cluster, SMA and premotor cortex showed the strongest similarity, followed by inferior parietal lobule and motor-somatosensory cortex. Interestingly, the closest pairs within each cluster (VV-LV in cluster 1, AI-IF in cluster 2, and SMA-PM in cluster 3), besides being anatomically close, also exhibited similar properties in the previous analyses (i.e., similar categories of preferred durations, similar local spatial associations between duration preferences, and similar degrees of clustering of duration preferences).

Overall, these findings suggest that changes in duration preferences along the cortical hierarchy are shaped by specific, non-trivial relationship between regions. When considered in light of our previous results, they point to a hierarchical organization of duration processing, where different areas showing different tuning properties may work together to support different outcomes of temporal processing. In occipital visual areas (cluster 1), duration is extracted from the sensory input and encoded through the monotonic increase in neuronal response amplitude. The two downstream clusters, instead, include parietal and frontal areas that support both duration readout and categorization. This suggests that these two clusters may contribute in parallel to different aspects of duration perception: one likely related to higher-level cognitive functions, involving the IPS, inferior frontal cortex, and anterior insula (cluster 2); and the other more motor-oriented, engaging the inferior parietal cortex and motor-related regions (cluster 3).

### Linking duration preferences to perception

Finally, we investigated whether and where, along the cortical hierarchy, duration preferences are linked to duration perception in the discrimination task. From each participant’s psychometric curve (shown in figure 7a), we derived the Point of Subjective Equality (PSE), defined as the comparison duration that is equally likely to be judged as longer or shorter than the reference duration. The PSE represents, therefore, the subjective boundary each participant employs to perform the task. We then computed the Kendall correlation between individual PSE values (*n* = 13) and median duration preferences of each individual stream and ROI (see *Methods - Analysis of the link between duration preferences and perception*). Figure 7 shows the correlation across streams (b) and ROIs (c).

**Figure 7:**
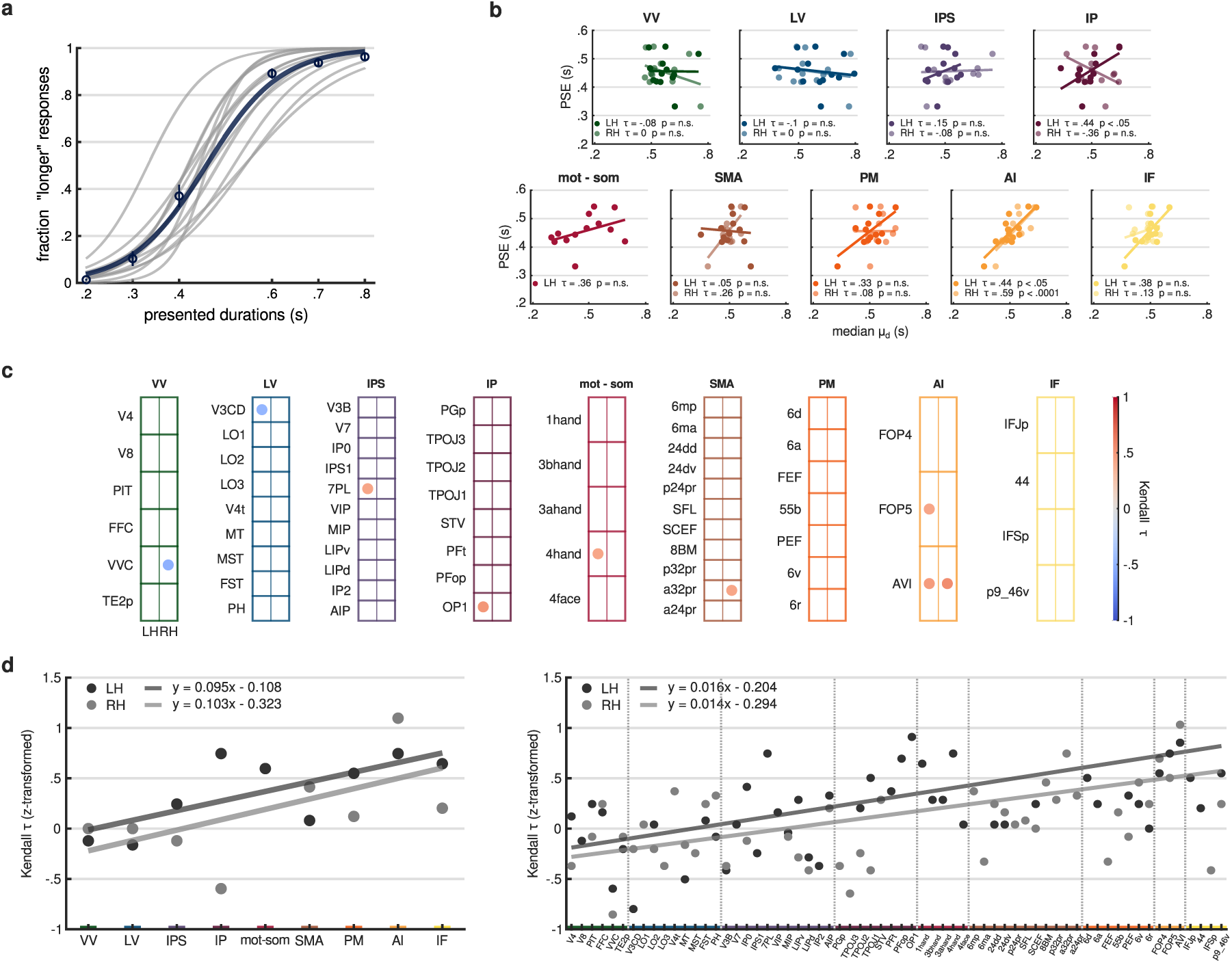
Correlation between PSE and duration preferences. **(a)** The plot shows individual psychometric curves in gray, with the group-level psychometric curve in blue. Circles represent the average fraction of “comparison longer than reference” responses across participants for each comparison duration, with error bars indicating standard errors. **(b)** Each scatter plot corresponds to a different stream, where individual PSE values (y-axis) are plotted against individual median preferred durations (x-axis) of that stream along with regression lines. Darker dots refer to the left hemisphere, while lighter dots refer to the right hemisphere. Kendall *τ* correlation values and their associated p-values are reported in the bottom left corner of each plot. **(c)** Each grid represents a different stream, with ROIs arranged in rows. The left (LH) and right (RH) hemispheres are displayed in separate columns. Each dot represents the Kendall correlation between individual PSE values and individual median preferred durations for a given ROI. Dot color indicates the Kendall *τ* value. Only significant (p-value *<* 0.05) correlations are shown. **(d)** Scatter plots represent z-transformed Kendall *τ* values across streams (left plot) and ROIs (right plot). Regression lines are displayed in gray, with their equations shown in the top left corner of each plot. Darker shade represents the left hemisphere, while lighter shade corresponds to the right hemisphere. In the right plot, vertical dashed lines separate different streams. In panels b, c, and d, the color code indicates different streams: green for VV, blue for LV, violet for IPS, purple for IP, red for mot-som, brown for SMA, orange for PM, ochre for AI, yellow for IF. See *Methods - Analysis of the link between duration preferences and perception*.

At the stream level, PSE values and median duration preferences were significantly and positively correlated in the left inferior parietal cortex (IP, *τ* = 0.44, p *<* 0.05) and in the anterior insula (AI), bilaterally (left: *τ* = 0.44, p *<* 0.05; right: *τ* = 0.59, p *<* 0.0001). When examining individual ROIs, we observed significant positive correlations in the left FOP5, belonging to AI (*τ* = 0.43, p *<* 0.05), in the bilateral AVI, belonging to AI (left: *τ* = 0.49, p *<* 0.05; right: *τ* = 0.56, p *<* 0.01), and in the right a32pr, belonging to SMA (*τ* = 0.43, p *<* 0.05). Interestingly, in these areas the majority of vertices showed, in our previous analyses, a preference for the medium duration range, which already suggested the representation of a categorical boundary used to perform the task. Finding a link between this type of duration preference and PSE further supports this idea and suggests that this representation has a subjective connotation, reflecting each participant’s individual perception in the discrimination task[31]. The other observed significant correlations are reported in figure 7c. Finally, to have a comprehensive view of how the correlation between PSE values and preferred durations changed along the cortical hierarchy, we fitted a regression line on z-transformed Kendall *τ* values, as shown in figure 7d, across streams (left plot) and ROIs (right plot). The correlation increased along the cortical hierarchy, shifting from values close to 0 in occipital ROIs to positive values in frontal ROIs (across left streams: *β* = 0.095, t(7) = 2.72, p *<* 0.05; across right streams: *β* = 0.103, t(6) = 1.49, p-value not significant; across left ROIs: *β* = 0.016, t(45) = 4.59, p *<* 0.0001; across right ROIs *β* = 0.014, t(44) = 3.59, p *<* 0.001). This last result suggests that, overall, the link between duration preferences and perception is progressively built up along the cortical hierarchy.

## Discussion

In this study, by means of a neuronal response model of unimodal duration tuning, we investigated how duration-tuned populations change across the cortical hierarchy, whether they are related across different areas, and whether they are linked to duration perception. We focused within cortical locations that showed significant activation at stimuli offset (i.e., when temporal information was fully available to participants), which corresponded to areas commonly reported in neuroimaging studies using explicit timing tasks of sub-second visual durations[2], [3].

Our findings indicate that, along the cortical hierarchy, neuronal populations show unimodal tuning to different ranges of durations. This is evident not only in their specific duration preferences, but also in their local spatial organization. In addition, these duration-tuned neuronal populations display different degrees of global spatial clustering across areas and are directly linked to duration perception only in specific regions. Specifically, the ventral and lateral visual areas mainly preferred long durations and displayed a lower degree of clustering compared to the other regions. The superior and inferior parietal cortex, lateral premotor cortex, and caudal SMA showed preferences spanning the full range of presented durations, with a higher degree of clustering. The anterior insula, inferior frontal cortex, and rostral SMA mainly preferred durations around the mean of the presented range, also displaying a higher degree of clustering. Notably, within specific areas of this latter group (a32pr within the SMA, and AVI and FOP5 within the anterior insula), duration preferences were positively correlated with PSE values. Finally, duration-tuned neuronal populations appeared to closely co-vary within different subsets of regions. This allowed us to identify three functional clusters: (i) occipital visual areas; (ii) intraparietal, inferior frontal, and insular areas; and (iii) inferior parietal, supplementary motor, premotor, and motor-somatosensory areas. Taken together, these findings suggest that distinct functional stages of duration processing are implemented in specific cortical regions, likely organized hierarchically and supporting different functions.

The first processing stage may occur in ventral and lateral visual areas, where we consistently observed a preference for long durations. This type of preference likely reflects a ramping response to stimulus duration, in line with previous findings showing that the BOLD response in early visual regions (from V1 to V3a, V3b and MT/V5) increases monotonically with stimulus duration[14], [15], [33]. The role of early sensory areas as gateways for time processing and perception has been suggested by diverse evidence, including psychophysics[34]–[36], brain imaging[15], [33], [37], [38], and brain stimulation[39], [40] works in humans as well as electrophysiology studies in rodents[30], [41]–[43]. In the tactile domain, for example, Reinartz and colleagues[43] showed that optogenetic excitation of the vibrissal somatosensory cortex in rats dilated perceived duration and amplified perceived intensity, suggesting that time perception is deeply rooted in sensory coding. In the visual domain, Tonoyan and colleagues[38] found that temporal biases induced in humans by temporal frequency adaptation could be predicted by the amplitude of an early ERP component (N200) recorded contralaterally to the stimulated visual field. This suggested that the subjective perception of time is linked to processes that begin locally and relatively early in the visual processing stream. In line with this latter study, a recent work from our group also showed that monotonic duration tuning in early visual areas is spatially dependent (i.e., only neuronal populations encoding the stimulus’s spatial position are modulated by duration)[15]. Collectively, these studies highlight the critical role of early sensory regions in duration processing and perception. Our findings further support this notion, suggesting that in the visual domain, monotonic responses may reflect an “accumulation” process of sensory information that encodes the duration of visual events[42]. This encoding mechanism could also account for the dilation of perceived time induced by the increase of low-level stimulus properties, such as speed[44], size[45], or visibility[46], by suggesting that a greater accumulation of sensory evidence leads to a longer encoded duration.

A second functional stage may occur in IPS and inferior parietal areas, lateral premotor regions, and caudal subdivisions of SMA. Here, we observed unimodal responses across the entire range of presented durations, well clustered in topographic maps[11]–[13]. At this stage, unimodal tuning and its topographic organization may provide the readout of temporal information, that could serve different purposes depending on the location along the cortical hierarchy[47]. On one hand, parietal maps may represent a relay stage, where temporal information from early visual areas is made available for further processing in frontal regions, consistently with the role of parietal cortex in sensory[48] and magnitude[49] binding. In the IPS, for example, different stimulus attributes such as duration, spatial position, size, and numerosity elicit topographically organized unimodal responses[50]. These responses not only coexist but also interact. Indeed, two recent fMRI studies from our group found that neuronal populations in the IPS selectively code for duration and spatial position[15], or duration and numerosity[13]. This suggests that duration readout may serve the integration of temporal information with other sensory information to create a unified stimulus representation. Furthermore, studies using transcranial magnetic stimulation showed the causal involvement of inferior parietal cortex in temporal judgments[39], [51], and an indirect measure of unimodal responses to durations in the right supramarginal gyrus has been shown to reflect perceived, rather than physical, durations[21]. These findings collectively suggest that unimodal tuning in parietal areas carries critical information for both sensory and perceptual accounts of temporal processing. On the other hand, temporal maps in lateral premotor areas and caudal SMA may support the readout of temporal information for task implementation and execution. In humans, these regions are widely activated in a variety of timing tasks[1]–[3]. In particular, the SMA is recognized as a core timing area[52], recruited regardless of duration range, stimulus modality, or task type. SMA activity depends also on the degree of attention directed towards the temporal attribute of the stimulus, with greater attention eliciting greater activity[53]. Indeed, unimodal neuronal responses to durations have been reported in SMA in presence of both perceptual and motor timing tasks, respectively in humans[11] and in monkeys[9], [10], but not when participants are only exposed to durations without engaging in judgments or tasks[12], [13]. Overall, these findings suggest that unimodal tuning in premotor cortices might be more oriented to behaviorally relevant processing.

A third functional stage may occur in the rostral SMA, anterior insula, and inferior frontal regions, where we observed well-clustered duration preferences around the mean of the presented temporal range. This finding aligns with previous fMRI studies employing passive viewing of changing durations, which showed that, in frontal regions, duration preferences either shift toward the mean of the tested range[13] or become more narrowly distributed[12]. These observations suggest that frontal areas process the overall statistics of the temporal environment and provide an abstract, categorical representation of durations, even when these durations are not immediately required for action. Our results extend this idea by also showing that in specific regions (i.e., a32pr in rostral SMA, FOP5 and AVI in anterior insula) duration preferences positively correlated with individual PSE values. This indicates that duration unimodal tuning at this stage may provide the representation of the perceptual temporal boundary used to solve the task. A similar finding was previously reported by Mendoza and colleagues[31], who found that pre-SMA neurons in monkeys performing an interval categorization task peaked near the boundary between short and long duration categories. A subgroup of these “boundary” neurons also adapted when the categorization criterion changed (i.e., when a different range of durations was tested). Crucially, the activity of these neurons reflected monkeys’ PSE values, rather than the physical boundary between categories, suggesting that they encode the subjective decision criterion that guides duration categorization behavior. Our findings suggest the presence of analogous “boundary” neuronal populations in humans, not only within the pre-SMA, but also in the anterior insula, a region associated with awareness, encompassing processes from interoception to perceptual decision-making[54]. Similar to the SMA, the anterior insula is also consistently implicated in timing tasks[3] and has been proposed to contribute to the subjective experience of duration by integrating interoceptive signals and emotional states over time[16]. Within this framework, Wittmann and colleagues observed that the activity in anterior insula, as well as in the inferior frontal cortex and pre-SMA, increased during the reproduction of supra-second durations, peaking just before the motor response terminating the reproduction[18]. This accumulator-like activity was suggested to reflect a temporal representation for an upcoming decision (i.e., stopping the reproduction), in line with the notion that the accumulation of bodily states and emotions underlies the perception of time[17]. While our experimental approach does not address the embodied aspects of temporal processing, our findings are consistent with the idea that frontal regions contribute to an internal and subjective representation of time. Importantly, we observed that this representation emerges from a cascade of transformations in neuronal tuning properties along the cortex. In addition, our findings suggest that, in the anterior insular cortex, boundary populations may constitute the neural substrate for integrating and directly comparing internal (e.g., bodily signals, temporal priors) and external (i.e., sensory) temporal information.

Finally, motor and somatosensory areas consistently showed preferences for short durations. However, all areas were restricted to the left hemisphere, consistent with the fact that participants provided their responses during the task using their right hand. This lateralized activation, combined with the preference for short durations, suggests that neuronal responses in these areas might reflect motor preparation rather than duration tuning. In each trial, the “ready” signal for motor response likely corresponded to the onset of each stimulus presentation, and among the tested durations, the one closer to the onset of the stimulus is the shortest one. In addition, similarly to occipital cortex, motor and somatosensory areas exhibited an overall weaker spatial autocorrelation compared to the other areas. This may further indicate the absence in these areas of unimodal responses for durations. This observation also strengthen the idea that spatial autocorrelation increases when efficient information processing and transmission are required[55], even when selectivity is restricted to a single duration, as we observed in more frontal areas.

An interesting finding concerns the response properties in the SMA, which are segregated along its rostro-caudal axis. Caudal regions showed unimodal responses to the full range of presented durations, while rostral regions showed preferences for the mean of the range. In addition, in one rostral region (a32pr), duration preferences also correlated with PSE values. The SMA is recognized as a heterogeneous structure, both anatomically and functionally, with a primary subdivision between SMA proper and pre-SMA, located caudal and rostral to the anterior commissure, respectively (for a review, see [56]). A functional rostro-caudal gradient has also been described for temporal processing[1], [57], [58]. For example, perceptual timing tasks are more likely to activate the pre-SMA, while motor timing tasks are associated with SMA proper[59]. Consistently with this view, the segregation we observed between tuning mechanisms may reflect this broader division of SMA, where a more categorical representation of durations (i.e., unimodal responses for the mean of the range) occurs in rostral areas, associated to more perceptual processing compared to caudal areas. Interestingly, the presence of different tuning mechanisms in the SMA has not been reported previously in humans, though it has been observed in monkeys[10], [31]. Overall, this suggests that the SMA may integrate two types of signals from different populations: one providing the readout of the duration of the stimulus at hand, and another representing the (subjective) criterion for solving a timing task. This integration could be a unique property of the SMA that makes it a core site for timing.

The study of long-range relationships between duration preferences revealed specific dependencies across brain areas, suggesting a hierarchical organization of the three functional stages described above - encoding, readout, and categorization. We identified three main functional modules where duration-tuned populations appeared to be more closely related: (i) occipital visual areas, (ii) intraparietal, inferior frontal, and insular areas, and (iii) inferior parietal, supplementary motor, premotor, and motor-somatosensory areas. The combination of their duration tuning properties with their involvement in other brain functions suggests that these modules may play different roles in temporal processing. The encoding stage involves occipital visual areas, which likely serve as the entry point for temporal information, where duration is extracted from incoming visual events. In contrast, the readout and categorization stages occur in the two downstream clusters, which engage distinct parietal and frontal areas. In one cluster, duration readout and categorization seem to function sequentially, with duration-tuned populations in the IPS conveying sensory information to boundary-populations in inferior frontal and anterior insular cortices. The IPS, working as a coordinate system for perception[60] with a central role in multisensory[61] and visuomotor processing[62], may integrate temporal information with other sensory inputs[13], [15]. This multimodal information may then be used by the inferior frontal and anterior insular cortices to generate a unified and flexible perceptual experience of the sensory environment. This cluster may therefore perform an abstraction of temporal information, potentially through the combination of areas that each implement distinct properties of duration tuning. Consistent with this idea of progressive abstraction, Hayashi and colleagues found that both the intraparietal and inferior frontal cortices store a common representation of temporal and numerical information. However, while the IPS processes this interaction at a perceptual level, the inferior frontal cortex represents it at a more abstract level[63]. The other cluster includes inferior parietal and motor-related regions, with almost all exhibiting neuronal populations that read out temporal information. The SMA is, in fact, the only area where both duration readout and categorization coexist. This cluster may be specialized in the motor implementation of timing decisions and behavior, and the discrete mapping of durations at hand may serve as a framework for immediate actions. The inferior parietal regions in this cluster, beyond their roles in visuospatial processing (PGp)[64] and visuomotor functions (PFop, PFt)[65], also contribute to higher-order cognitive processes such as memory retrieval (PGp)[65], theory of mind (TPOJ1)[66], and emotional regulation (STV)[67] (see also [29], [68]). Moreover, areas of the inferior parietal lobule have been causally linked to duration perception[21], [39], [51], [69]. These areas may thus represent temporal information in a behaviorally relevant manner and relay it to downstream motor-related areas, where timing behavior is ultimately implemented, initiated, and controlled.

It is important to note that we limited our investigations exclusively on cortical regions, without considering contributions from the cerebellum and subcortical structures, which are known to play a critical role in temporal processing[70]–[75]. In addition, we only focused on the visual modality, and the extent to which these findings may generalize to other sensory modalities remains uncertain. For example, in the auditory domain, duration tuning have been found in primary auditory cortices only[76], and a recent review identified different activation patterns for auditory and visual timing[3].

Overall, this study successfully replicates, with higher anatomical precision, classical findings from fMRI meta-analyses on the localization of temporal processing, while directly linking them with previous electrophysiology and neuroimaging results regarding the underlying neural mechanisms. This work proposes a functional cortical hierarchy for visual temporal processing, where different areas contribute differently to duration perception, as reflected in their neuronal tuning properties. Additionally, it sheds light on the neural basis of the subjective experience of time, suggesting that this may be rooted in the fundamental tuning properties of brain responses.

## Methods

The data presented in this work have been previously used for a different publication[15].

### Participants

Thirteen healthy volunteers participated in this study (6 females; sex information self-reported; gender information not collected; mean age = 29.6, SD = 7.3; 2 left-handed participants). All volunteers had normal or corrected-to-normal visual acuity. The experimental procedures were approved by the International School for Advanced Studies (SISSA) ethics committee (protocol number 11773) in accordance with the Declaration of Helsinki. All participants gave their written informed consent to participate in the experiment and they were financially compensated for their time and travel expenses.

### Stimuli and procedure

#### Stimuli

Participants were presented with visual stimuli displayed on a BOLD screen (Cambridge Research Systems 32-inch LCD widescreen, resolution = 1920 × 1080 pixels, refresh rate = 120 Hz) placed at a total viewing distance of 210 cm outside the scanner bore and viewed via a mirror. The stimuli were colored circular patches of Gaussian noise subtending 1.5^◦^ of visual angle, changing dynamically frame by frame and presented on a gray background. Each stimulus was constructed by randomly selecting RGB values from a Gaussian distribution of mean = 127 and SD = 35 for each of its pixels and frames. This ensured that the average stimulus luminance was constant and independent of its duration. To prevent the perception of flickers induced by the fast-changing rate in the stimulus, we scaled down its pixel resolution (scaling factor = 12.33). This created a blurring effect that homogenized the local contrasts of the stimulus over frames and minimized possible flickering effects[77]. The entire experimental procedure was generated and delivered using MATLAB and Psychtoolbox-3[78]. An identical set-up was used during participants training.

#### Task and experimental design

Participants were asked to perform a single interval discrimination task. They had to compare the duration (i.e., display time) of a comparison stimulus to the duration of a reference stimulus internalized during the training procedure. The task was to report whether the comparison stimulus was longer or shorter than the reference. The reference duration was 0.5 s, and the comparison durations were 0.2, 0.3, 0.4, 0.6, 0.7, and 0.8 s. The comparisons were presented in different spatial positions (i.e., display locations on the screen), which could be either at 0.9^◦^ or 2.5^◦^ of visual angle diagonally from the center of the screen in the lower-left or lower-right visual quadrant (see the close-up in figure 1a). Stimuli did not overlap across spatial positions. A white fixation cross (0.32^◦^ of visual angle) was displayed at the center of the screen throughout the experiment. In each trial, the comparison stimulus was presented in a specific position and entailed a specific duration. After a randomized interval from the offset of the stimulus (stimulus-cue interval (SCI) uniformly distributed between 0.9 and 1.2 s), the participants’ response was cued with a color switch (from white to black) in the fixation cross. The response was allowed within a 2 s window, but no emphasis was placed on reaction times. Participants were instructed to provide their responses by pressing one of two buttons on a response pad with their right index finger or right middle finger to express the choices “comparison longer than reference” and “comparison shorter than reference” respectively. No feedback was provided after the response. A uniformly distributed inter-trial interval (ITI) between 1.8 and 2.5 s interleaved the trials. See figure 1a for a pictorial representation of the trial structure. The stimulus duration varied trial by trial in a pseudo-randomized and counterbalanced fashion, whereas its position varied sequentially and cyclically to minimize attentional switching effects on the duration judgment[79]. Each cycle started and ended at 2.5^◦^ in the lower-left quadrant and comprised a clockwise and counterclockwise presentation of the stimulus in all positions, from 2.5^◦^ lower-left to 2.5^◦^ lower-right and back (i.e., 2.5^◦^ L, 0.9^◦^ L, 0.9^◦^ R, 2.5^◦^ R, 2.5^◦^ R, 0.9^◦^ R, 0.9^◦^ L, 2.5^◦^ L). Each half cycle (i.e., when the presentation order turned from clockwise to counterclockwise and vice versa) was followed by a 2.64 s (2 TR) interval. To ensure a balanced presentation of all combinations of durations and positions within each block, a cycle was repeated six times, and each duration was presented twice in each position, for a total of 48 trials per block. Each participant performed 10 blocks inside the scanner acquired in separate fMRI runs. Duration randomization differed in each block, whereas the position sequence was always the same. Stimuli presentation was synchronized with the scanner acquisition at the beginning and at the middle of each cycle. Participants were instructed to maintain their gaze at the fixation cross while performing the task, and eye movements were recorded with an MR-compatible eye-tracking system (R Research Eyelink 1000 Plus) placed inside the scanner bore.

#### Training

Participants underwent a training procedure outside the scanner to familiarize with the stimuli and the task. First, they were asked to internalize the duration of the reference stimulus. In this phase, participants passively viewed a 0.5 s stimulus presented at the center of the screen three times (inter-stimulus interval uniformly distributed between 1.8 and 2.5 s) in each trial. They were free to complete as many trials as they needed to feel confident they had internalized the duration of the stimulus. Next, participants performed the first training block of the duration discrimination task. The task structure was identical as described before (see *Methods - Task and experimental design*), but all comparisons were presented at the center of the screen. This was done to ensure that participants were able to correctly discriminate the comparisons from the reference stimulus. Finally, participants performed a second training block identical to the experimental blocks to familiarize with the experimental procedure. Throughout the training phase, participants received visual feedback about their performance and eye movements.

### MRI acquisition

MRI data were acquired with a Philips Achieva 7T scanner equipped with an 8Tx/32Rx-channel Nova Medical head coil. T2*-weighted functional images were acquired using a three-dimensional EPI sequence with anterior-posterior phase encoding direction and the following parameters: voxel resolution = 1.8 mm isometric; repetition time (TR) = 1.32 s; echo time (TE) = 0.017 s; flip angle = 13 degrees; bandwidth = 1750 Hz/px. Universal kt-points pulses were used to achieve a more homogeneous flip angle throughout the brain[80]. The matrix size was 112×112×98, resulting in a field of view of 200(AP) x 200(FH) x 176.4(LR) mm. At the end of each run, 4 volumes were acquired with the opposite phase encoding direction in order to perform susceptibility distortion correction (see *Methods - MRI data preprocessing*). A minimum of 190 volumes (acquisition time ≈ 4 minutes) was acquired for each experimental run. Peripheral pulse and respiratory signals were recorded simultaneously with the fMRI data acquisition using the Philips MR Physiology recording system. The finger clip of the peripheral pulse unit was placed on the subject’s left ring finger, and the respiratory sensor was placed over the diaphragm and secured with a band. Eye movements were monitored and recorded with an eye-tracking system (SR Research Eyelink 1000 Plus) mounted onto a hot-mirror system and located inside the scanner bore. High-resolution T1-weighted images were obtained using the MP2RAGE pulse sequence[81] optimized for 7T (voxel size = 0.7×0.7×0.7 mm, matrix size = 352×352×263).

### MRI data preprocessing

Pulse-oximetry and respiratory components were regressed out from functional MRI data before the preprocessing. We converted physiological signals into slice-based regressors using RetroTS.py (AFNI), and we performed the Retrospective Image Correction with a custom routine based on 3dretroicor (AFNI). This procedure was applied only when physiological signals showed a reliable frequency spectrum (they were removed in 100 out of 130 runs). MRI data were preprocessed using fMRIPrep 21.0.2[82], [83], which is based on Nipype 1.6.1[84], [85]. See the supplementary methods for details about the pipeline.

### Behavioral performance analysis

We first assessed that the display location of the stimulus did not bias duration discrimination or affect its sensitivity (see [15] for full statistical analysis and results). Next, we estimated individual psychometric curves based on the average fraction of “comparison longer than reference” responses for each comparison duration, collapsing across spatial positions (figure 7a, gray curves). We also computed the mean fraction of “comparison longer than reference” responses across participants for each comparison duration to estimate the group psychometric curve shown in blue in figure 7a. All psychometric curves were fitted using the glmfit function in MATLAB with a logit link function. For each participant, we then derived the Point of Subjective Equality (PSE), which represents the comparison duration equally likely to be judged as longer or shorter than the reference and thus quantifies the bias in the duration discrimination task. Individual PSE values were used to investigate the link between duration tuning and perception as described in *Methods - Analysis of the link between duration preferences and perception*.

### General linear model (GLM) analysis

We initially analyzed functional MRI data, resampled on the cortical native surface (Freesurfer’s fsnative), with a General Linear Model (GLM) approach using the GLMdenoise toolbox[86]. For each run, the design matrix included one regressor for each combination of comparison duration and position, time-locked to the offset of each stimulus (events of interest), and one regressor time-locked to the onset of each response (event of no interest). In total, we modeled 25 events (6 stimulus durations x 4 stimulus positions + response). As GLMdenoise automatically estimates noise regressors, no motion correction parameters were entered in the procedure. Regressors were convolved with the canonical hemodynamic response function. For each subject, this procedure yielded a set of 100 bootstrapped beta weights for each vertex. In the subsequent modeling procedure, we used the median beta weights across bootstraps, converted to percentage of signal change.

### Population receptive field (pRF) modeling

To estimate the tuning properties of BOLD responses elicited by stimulus duration, we modeled the set of GLM beta weights (see *Methods - General linear model (GLM) analysis*) using the Population Receptive Field (pRF) approach[27]. We applied a model that assumes a Gaussian response to stimulus duration and invariance to stimulus spatial position. The model is described by the following equation:

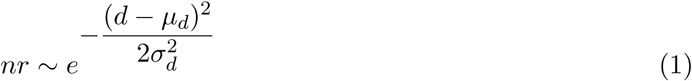

where *µ_d_* represents the duration preference (i.e., the stimulus duration eliciting the greatest neuronal response) and *σ_d_* denotes the sensitivity of the response to changes in stimulus duration (see figure 1b).

#### Fitting procedure

We first translated our experimental manipulations of the stimulus in a three-dimensional matrix. The stimulus spatial and temporal dimensions were mapped along the first and second axes in arbitrary units ranging from 1 to 100. The third dimension represented all combinations of stimulus duration and position, ordered according to GLM beta weights. To derive the predicted neuronal response, we multiplied the pRF model by the three-dimensional stimulus matrix and then integrated over the first and second dimensions, yielding a series of 24 predictions (i.e., one for each combination of stimulus duration and position). For each vertex of the cortical surface, we optimized this predicted neuronal response by minimizing the residual sum of squares relative to the GLM beta weights. This optimization process was performed in two steps. First, a grid search tested the performance of a large set of parameters for the pRF model. Next, the best-performing parameters from the grid fit were used as seed in an iterative procedure that explored previously untested parameter combinations. The iterative optimization was based on the Nelder-Mead method[87], as implemented in the fminsearchbnd function in MATLAB. Only parameters accounting for at least 10% of variance in the grid fit were further optimized in the iterative step. Vertices displaying a negative pRF were excluded from further analyses. We performed the entire procedure using custom functions in MATLAB. Figure 1b shows the fitting result for a representative vertex.

### Regions of interest (ROIs) identification

We conducted our analyses within a set of ROIs identified using a mass-univariate GLM approach performed with the Statistical Parametric Mapping toolbox in MATLAB (SPM12, version 7219, Wellcome Department of Imaging Neuroscience, University College London). The volumetric functional data, resampled on the MNI152NLin6Asym space, were first smoothed with a 2 mm FWHM Gaussian kernel. For the first-level analysis, we constructed a design matrix that included one regressor for each combination of comparison duration and position, time-locked to the offset of the stimuli (events of interest), and one regressor time-locked to the onset of the responses (event of no interest). All event durations were set to zero, and regressors were convolved with the canonical hemodynamic response function. The model also included 6 motion correction parameters. For each subject, we then estimated one contrast for each stimulus duration, regardless of spatial position. The resulting contrast images were entered into a second-level full-factorial analysis, where an overall t-contrast was computed. The statistical threshold was set at p*<*0.001, FWE cluster-level corrected for multiple comparisons across the entire brain volume, with cluster size estimated at p FWE-uncorrected = 0.001. The volume of group-level clusters was then resampled onto the fsaverage surface using FreeSurfer’s mri vol2surf. To minimize potential statistical inflation due to surface resampling, we applied a minimum t-value threshold of 4. Additionally, small clusters were excluded using a minimum surface threshold of 20 *mm*^2^.

To identify the anatomical locations of t-value clusters, we used the HCP MMP 1.0 atlas[25] in fsaverage space. An atlas area was considered an ROI if at least 5% of its surface was covered by one cluster. For clusters within motor and somatosensory areas, we used the same procedure using the topological atlas by Sereno and colleagues[26] to achieve finer parcellation. This process resulted in the identification of 47 ROIs in the left hemisphere and 46 ROIs in the right, with 29 ROIs shared between hemispheres. Finally, we grouped ROIs into nine functional streams based on their anatomical locations and their description provided in the multipart supplement of the HCP atlas[88]. All ROIs are listed in table 1 and displayed in figure 1c. After identification on the fsaverage surface, the ROIs were subsequently extracted from each participant’s native surface for further analyses.

**Table 1:** Regions of interest (ROIs). ROIs are listed separately for the left and right hemispheres and grouped by functional streams. The ROI nomenclature follows that provided with the atlases[25], [26].

| Functional stream | Left hemisphere ROIs | Right hemisphere ROIs |
| --- | --- | --- |
| Ventral visual (VV) | V4, V8, PIT, FFC, VVC, TE2p | V4, PIT, FFC, VVC, TE2p |
| Lateral visual (LV) | V3CD, LO2, MT, MST, FST, PH | V3CD, LO1, LO2, LO3, V4t, MT, MST, FST, PH |
| Intraparietal sulcus (IPS) | V3B, V7, IP0, IPS1, 7PL, VIP, MIP, LIPv, LIPd, IP2, AIP | V3B, IP0, MIP, LIPv, LIPd, AIP |
| Inferior parietal (IP) | TPOJ2, TPOJ1, PFt, PFop, OP1 | PGp, TPOJ3, TPOJ2, TPOJ1, STV |
| Motor-somatosensory (mot-som) | 1-hand, 3b-hand, 3a-hand, 4-hand, 4-face |  |
| Supplementary motor areas (SMA) | 24dd, 24dv, SCEF | 6mp, 6ma, 24dd, 24dv, p24pr, SFL, SCEF, 8BM, p32pr, a32pr, a24pr |
| Premotor (PM) | 6d, 6a, PEF, 6v, 6r | FEF, 55b, PEF, 6v, 6r |
| Anterior insula (AI) | FOP4, FOP5, AVI | FOP4, FOP5, AVI |
| Inferior frontal (IF) | IFJp, 44, p9_46v | IFSp, p9_46v |

### Analysis of duration preference changes along the cortical hierarchy

In this first analysis, we investigated how the distribution of duration preferences (i.e., *µ_d_* parameters) changed across different areas. For each participant, we calculated the median *µ_d_* within each ROI, including both left and right hemispheres for bilateral ROIs. We then tested two linear mixed effect (LME) models, using the lme4 R package[89], with the following formulas:

To test changes across streams:

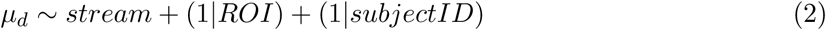

To test changes across ROIs:

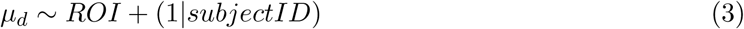

The variance explained by each LME model was computed using the MuMIn package[90]. The Satterwaite’s method[91] implemented in the lmer package[92] was used to estimate the degrees of freedom for the LME model ANOVA. Estimated marginal means were compared using the emmeans package[93], which employs the Kenward-Roger’s method[94] for degrees of freedom estimation.

### Analysis of duration preference categories along the cortical hierarchy

We further characterized changes in duration preferences along the cortical hierarchy by performing a categorization analysis in conjunction with the local Moran’s I statistic.

#### Categorization of duration preferences

For each participant, we categorized vertices based on their duration preference into five groups: short (.2-.32 s), mid-short (.32-.44 s), medium (.44-.56 s), mid-long (.56-.68 s), and long (.68-.8 s). We then computed the fraction of vertices in each category for each stream and ROI, averaging across hemispheres for bilateral ones. These data were analyzed using two separate two-way repeated-measure ANOVA, with category and either stream or ROI as within-subject factors, using the anova test function in R (from the rstatix package[95]). Marginal means were estimated using the lm function in R, from the stats package[96], with the following model formulas: 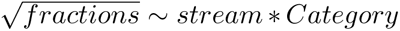 for streams and 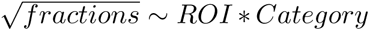 for ROIs. Multiple comparisons were performed with the emmeans package[93]. The square-root transformation improved the normality of the data, ensuring more robust contrast estimates.

#### Local Moran’s I

For each vertex of the cortical surface, we computed the local Moran’s I statistic[97], which quantifies the correlation between a vertex’s duration preference and the average preference of its neighboring vertices, along with an associated significance assessment. This analysis allowed us to identify spatial clusters of significantly associated vertices and classify their spatial relationships into four types: high duration preference surrounded by high (high-high, or HH); low surrounded by low (low-low, or LL); high surrounded by low (high-low, or HL); and low surrounded by high (low-high, or LH). For each participant, we computed the local Moran’s I across the entire cortical surface of each hemisphere separately, centering duration preferences to the mean preference of that hemisphere. The neighborhood structure included the same number of vertices for all ROIs within a given hemisphere and participant, corresponding to one-fourth of the vertices in the smallest ROI of that hemisphere. Overall, the neighborhood structure included between 59 and 135 vertices. This approach allowed us to capture broad spatial associations in duration preferences and compare them across ROIs. To validate the statistic, we used a conditional permutation procedure in which we computed the statistic 999 times. In each iteration, duration preferences were randomly shuffled among neighboring vertices while keeping the value of the current vertex fixed. Vertices with a p-value below 0.05 were considered significant. We implemented this statistic with custom MATLAB functions, based on the GeoDa software[98]. Once we obtained the local cluster map for each hemisphere and participant, we computed the fraction of vertices assigned to each spatial association type (i.e., LL, HH, LH, HL) within each ROI. Since HL and LH associations were nearly absent (see supplementary figure 1), our subsequent statistical analysis focused on the difference in the fraction of vertices between LL and HH. The LL-HH difference was computed for each participant within each ROI and averaged across hemispheres for bilateral ROIs. We then analyzed these data using the following LME models:

To test changes across streams:

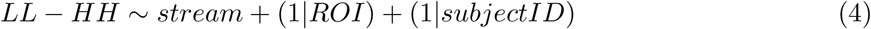

To test changes across ROIs:

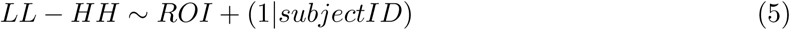

This analysis was performed using the same tools described previously (see *Methods - Analysis of duration preference changes along the cortical hierarchy*). Finally, to examine the overall trend of LL-HH differences across the cortical hierarchy, we averaged them across participants within each ROI and fitted a linear regression line (using the fitlm function in MATLAB) to the data ordered from occipital to frontal regions, as shown in figures 4b and 4d. For changes across streams (panel b), LL-HH differences were also averaged across ROIs belonging to the same functional stream. We then extracted the slope coefficient with its associated t statistics. To further validate the previous analyses, we also tested the LL and HH fraction values. We conducted two separate two-way repeated-measure ANOVA, with association type (i.e., LL and HH) and either stream or ROI as within-subject factors, using the anova test function in R (from the rstatix package[95]). Marginal means were estimated using the lm function in R, from the stats package[96], with the following model formulas: *fractions* ∼ *stream* ∗ *Type* for streams and *fractions* ∼ *ROI* ∗ *Type* for ROIs. Multiple comparisons were performed with the emmeans package[93]. We verified that the estimated marginal means and contrasts were numerically identical to those obtained from LME models including a random intercept for each subject, which produced a singular fit in some cases. The results are reported in supplementary tables 15 - 20.

### Analysis of the topographic organization of duration preferences along the cortical hierarchy

We assessed the topographic properties of duration preferences across the cortical hierarchy with two spatial statistics methods: the global Moran’s I statistic and the experimental variogram.

#### Global Moran’s I

The global Moran’s I statistic assesses the presence of clustering within a sample by quantifying its overall spatial autocorrelation. It is defined as the average of all local Moran’s I statistics (i.e., the correlation between each vertex’s duration preference and the average preference of its neighbors). A Moran’s I value close to 0 indicates a random spatial distribution of duration preferences, while values above or below 0 indicate increasing degrees of spatial clustering. A positive Moran’s I indicates that vertices with similar duration preferences are spatially close, whereas a negative Moran’s I reflects an inverse relationship, where vertices with high duration preferences are near those with low duration preferences, or vice versa. We computed this statistic separately for each ROI, centering duration preferences to the mean preference of that ROI. The neighborhood structure was defined using the 12 nearest neighboring vertices for each vertex. To estimate p-values, we performed a random permutation test, shuffling preferred durations across vertices and recomputing the statistic 999 times. Because each ROI has a unique neighborhood structure, statistical comparisons of Moran’s I values across ROIs are not feasible. To obtain the global Moran’s I for each functional stream, we averaged the statistic across the corresponding ROIs.

#### Variogram

The experimental variogram quantifies the spatial autocorrelation in a sample by measuring how the variance between pairs of vertices changes as a function of their distance. For each ROI, we constructed the variogram by first computing the pairwise distances between all vertices within the region. These distances were then grouped into 2mm-resolution bins, and for each bin, we calculated the variance in duration preferences across vertices. This approach ensured that variance estimation remained independent of the extent of the ROI. Duration preferences of each ROI were z-scored prior to variogram computation. We implemented this procedure using custom MATLAB functions based on the GeostatsPy package[99]. From each variogram, we then extracted two parameters, the nugget and the range. The nugget represents the variance at the shortest distance between vertices (see the inset in figure 5b and supplementary figure 3a). The range corresponds to the distance at which the total variance of the ROI, calculated without accounting for spatial structure, is first reached (see the inset in figure 5c and supplementary figure 4a). We then studied how nuggets and ranges varied across both functional streams and ROIs using the following LME models:

To test changes across streams:

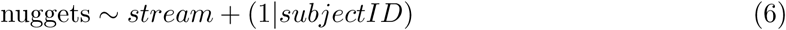

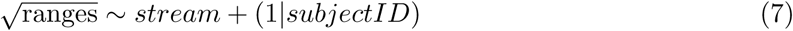

For this analysis, nuggets and ranges were averaged across hemispheres and ROIs within each functional stream.

To test changes across ROIs:

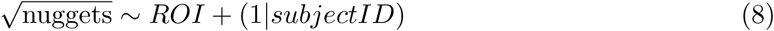

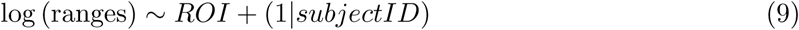

For this analysis, nuggets and ranges were averaged across hemispheres for bilateral ROIs.

These analyses were performed using the same tools described previously (see *Methods - Analysis of duration preference changes along the cortical hierarchy*). Square-root and log transformations were used to improve normality in the sample data and ensure more robust model performance.

### Analysis of long-range relationships between duration preferences

We investigated whether and how changes in duration preferences are related along the cortical hierarchy by performing a correlation analysis across areas. For each participant, we computed the median duration preference within each ROI, including both hemispheres for bilateral ROIs. We then calculated the Kendall correlation matrix across areas using the cor function from the R stats package[96]. To gain a more generalized view of these relationships, we repeated this computation using the median duration preferences of each functional stream per participant, again including both hemispheres where applicable. This correlation matrix was then used to construct a dendrogram. First, we converted the correlation matrix into a distance matrix using the dist function. Hierarchical clustering was then performed with the hclust function (using the default “complete” method), and the dendrogram was generated using the as.dendrogram function (all functions are from the R stats package[96]).

### Analysis of the link between duration preferences and perception

We investigated whether and where a link exists between duration tuning and perception by performing a correlation analysis between duration preferences and PSE values (see *Methods - Behavioral performance analysis*). For each participant, we computed the median duration preference within each functional stream and ROI. We then calculated the Kendall correlation within each stream and ROI between individual duration preferences and PSE values, using the corr function in MATLAB. To examine the trend of this correlation along the cortical hierarchy, Kendall’s *τ* values were transformed to z values using the formula 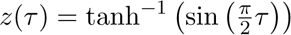 [100], ordered from occipital to frontal regions (as shown in figure 7d), and fitted with a linear regression line using the fitlm function in MATLAB. Finally, we extracted the slope coefficient with its associated t statistics.

## Acknowledgments

We thank the staff of the Spinoza Centre for Neuroimaging (www.spinozacentre.nl) for their help and assistance during data collection. This project has received funding from the European Research Council (ERC) under the European Union’s Horizon 2020 research and innovation programme (grant agreement no. 682117 BIT-ERC-2015-CoG) to D.B., and from the Italian Ministry of University and Research under the call FARE (project ID: R16X32NALR) and under the call PRIN2017 (project ID: XBJN4F) to D.B.

## Author Contributions Statement

V.C. and G.F.: conceptualization, data curation, formal analysis, investigation, methodology, software, validation, visualization, writing - original draft, writing - review & editing; D.B.: conceptualization, funding acquisition, project administration, resources, supervision, writing - review & editing.

## Competing Interests Statement

The authors declare no competing interests.

## Data Availability

The raw MRI data are protected and are not available due to data privacy laws. All processed data supporting our findings will be available upon publication in OSF at the following link (now private): https://osf.io/2tequ/.

## Code Availability

Codes supporting our findings will be available upon publication in OSF at the following link (now private): https://osf.io/2tequ/.

## Notes

### Competing Interest Statement

The authors have declared no competing interest.

### Summary of Updates

Results of local Moran's I statistic (Results - Categorization of duration preferences along the cortical hierarchy; Methods - Analysis of duration preference categories along the cortical hierarchy - Local Moran's I; Figures 4b and 4d; Supplementary Figure 1; Supplementary Tables 13 - 20) Order of SMA regions (Figures 3, 4, 6, 7; linear fit in Results - Linking duration preferences to perception)

