## Supplementary Materials for "Defining a functional hierarchy of millisecond time: from visual stimulus processing to duration perception"

### Supplementary Figures

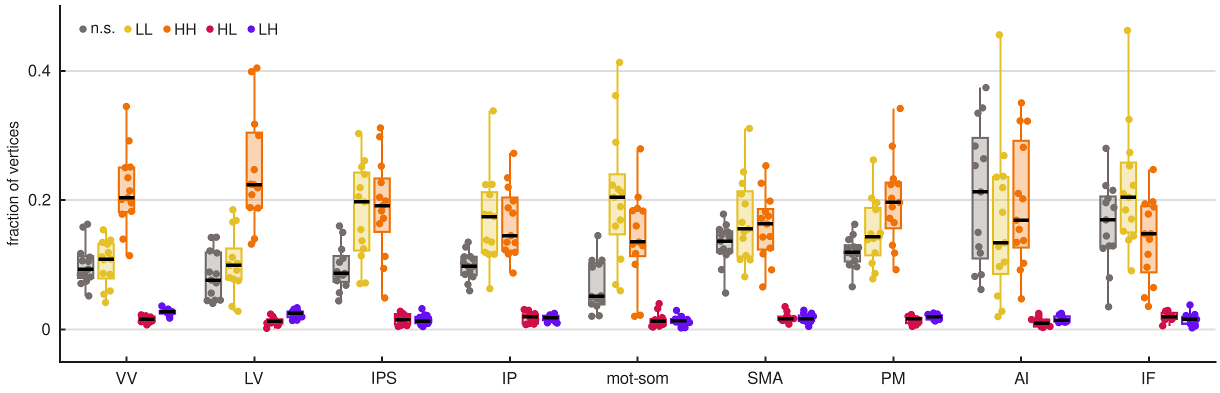


Supplementary figure 1: **Local spatial associations between duration preferences.** Box plots show the group-level distribution (n = 13) of the individual fraction of vertices assigned to each type of local spatial association (based on Moran's I statistic) across the different functional streams. Spatial association types are color-coded: gray for non significant (n.s.), yellow for low-low (LL), orange for high-high (HH), pink for high-low (HL), and purple for low-high (LH). Each dot represents the mean fraction across ROIs and hemispheres for each participant. The horizontal black line indicates the median of the distribution, the box shows the interquartile range, and whiskers represent the minimum and maximum values. Streams are color-coded on the x-axis: green for ventral visual areas (VV), blue for lateral visual areas (LV), violet for intraparietal sulcus (IPS), purple for inferior parietal areas (IP), red for motor and somatosensory areas (mot-som), brown for supplementary motor areas (SMA), orange for premotor areas (PM), ochre for anterior insula (AI), yellow for inferior frontal areas (IF). See *Methods - Analysis of duration preference categories along the cortical hierarchy - Local Moran's I.*

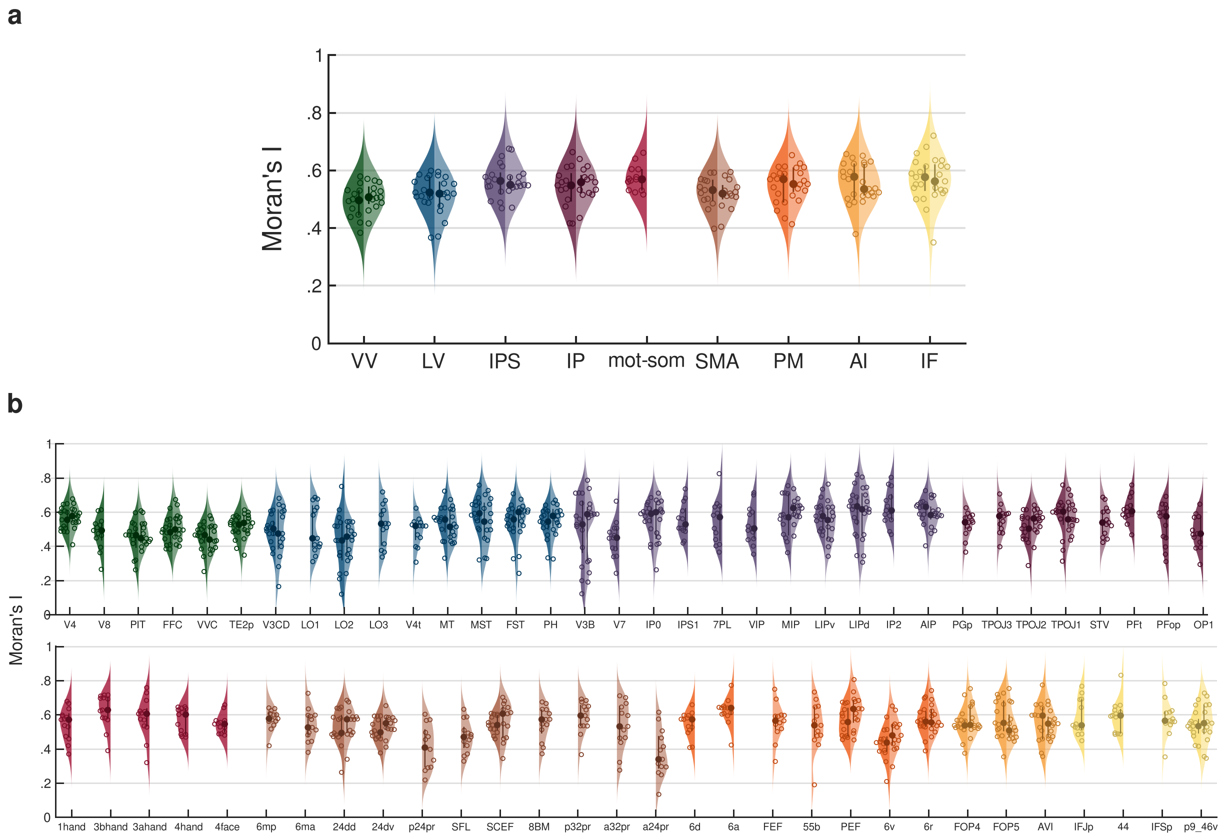


Supplementary figure 2: **Group-level distributions of Moran's I statistic values.** Each violin plot represents the group-level distribution (n = 13) of Moran's I statistic values across streams **(a)** and ROIs **(b)**. Both streams and ROIs are ordered from occipital to frontal and from dorsal to ventral. Streams are color-coded as follows: green for VV, blue for LV, violet for IPS, purple for IP, red for mot-som, brown for SMA, orange for PM, ochre for AI, yellow for IF. ROIs are color-coded according to their respective streams. The left side of each violin represents the left hemisphere (darker shades), while the right side represents the right hemisphere (lighter shades). Dots indicate the median of each distribution, while circles correspond to individual data points. Thick lines represent interquartile ranges. The kernel density estimates were computed using a 7% bandwidth. See *Methods - Analysis of the topographic organization of duration preferences along the cortical hierarchy - Global Moran's I*.

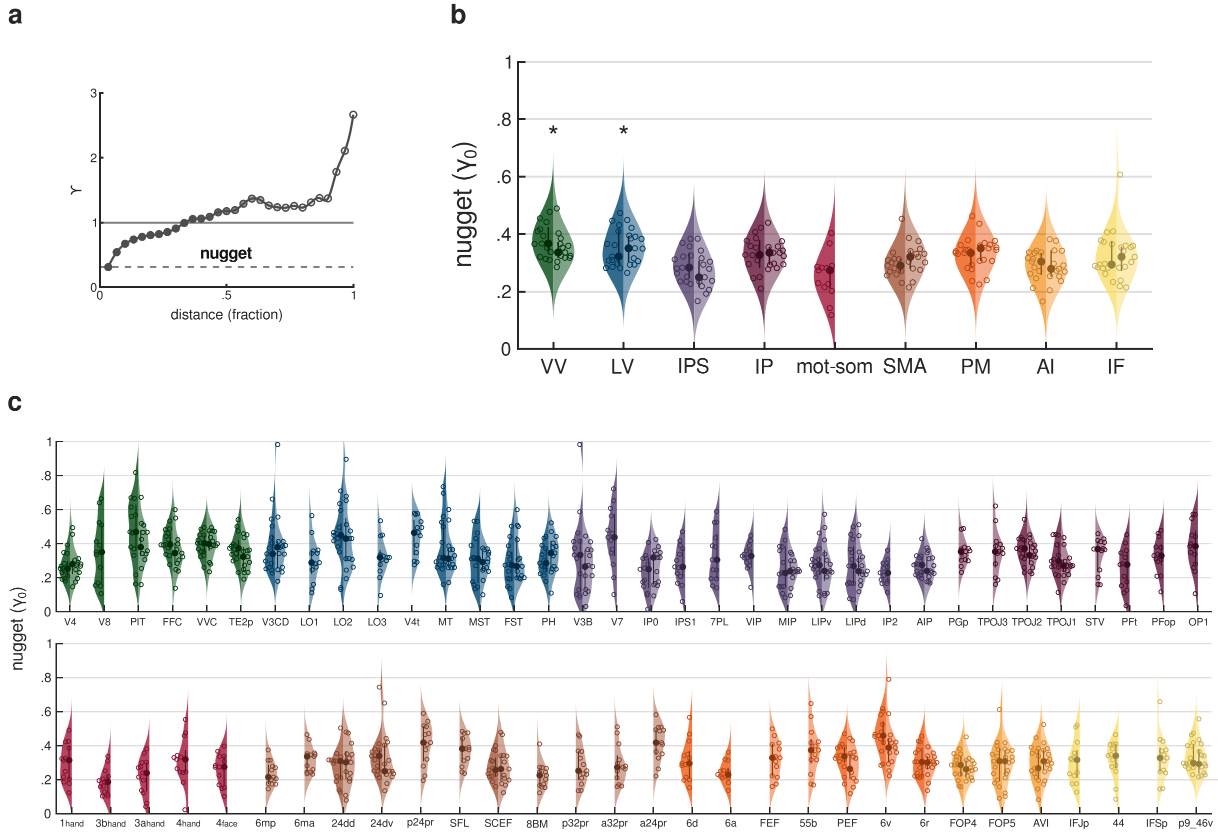


Supplementary figure 3: **Group-level distributions of nuggets.** **(a)** An example variogram highlighting the nugget is shown. Each dot represents the variance ($\gamma$) in duration preferences between pairs of vertices at increasing distance (expressed as a fraction of the total extent of the ROI). Dots shading reflects the number of vertex pairs at each distance, with darker shades indicating a higher count. The solid line ($\gamma = 1$) indicates the total variance of the ROI, computed without accounting for spatial structure. The nugget (i.e., the variance at the shortest distance) is highlighted by the dashed line. Panels **(b)** and **(c)** show the group-level distributions ($n = 13$) of variogram nuggets across streams and ROIs, respectively. Nugget values are expressed as a fraction of the total variance of the ROI. Graphical details are the same as in supplementary figure 1. Asterisks in panel b indicate streams that are statistically different from the others (see supplementary table 23). See *Methods - Analysis of the topographic organization of duration preferences along the cortical hierarchy - Variogram.*

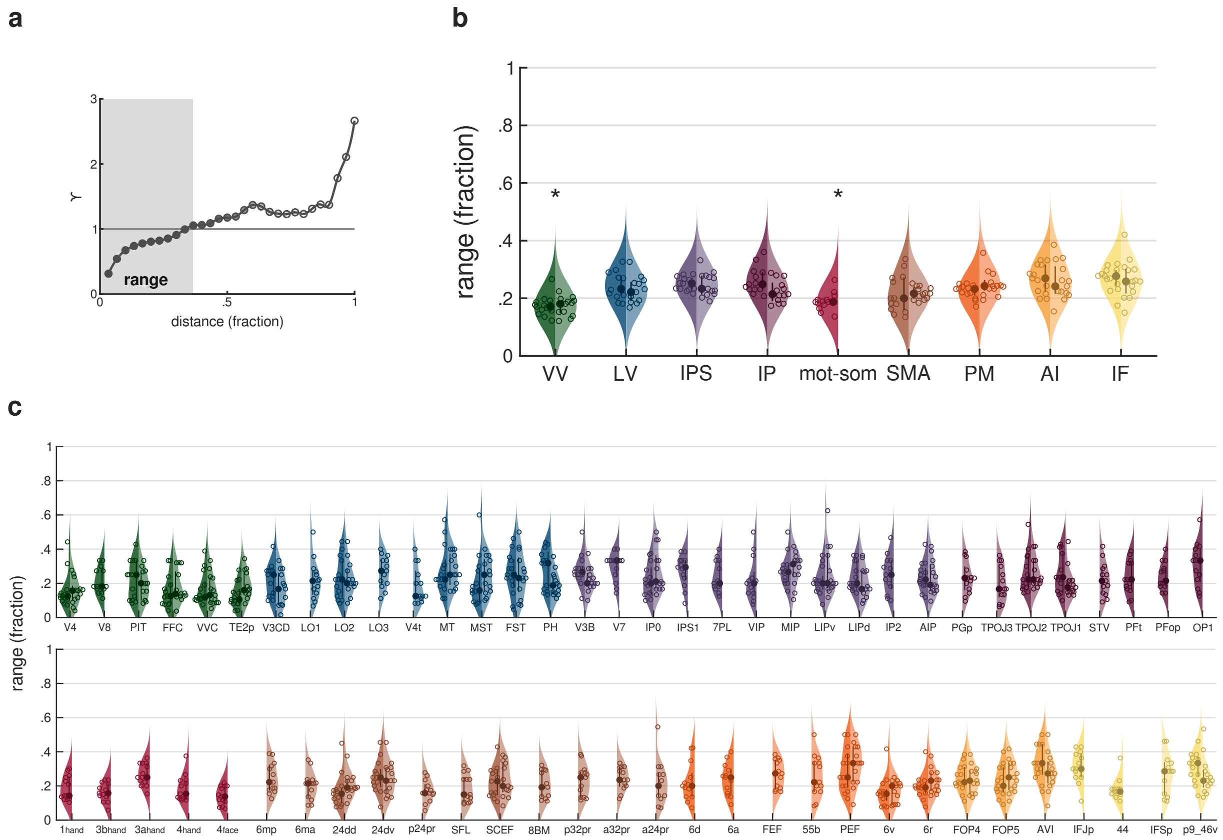


Supplementary figure 4: **Group-level distributions of ranges.** **(a)** The same example variogram from supplementary figure 2 is shown, with the range (i.e., the distance at which the total variance of the ROI is reached) highlighted by a gray box. Panels **(b)** and **(c)** show the group-level distributions (n = 13) of variogram ranges across streams and ROIs, respectively. Range values are expressed as a fraction of the maximum distance between vertices of the ROI. Graphical details are the same as in supplementary figure 1. Asterisks in panel b indicate streams that are statistically different from the others (see supplementary table 24). See *Methods - Analysis of the topographic organization of duration preferences along the cortical hierarchy - Variogram.*

### Supplementary Tables

| | Sum.Sq | Mean.Sq | NumDF | DenDF | F.value | Pr(>F) |
|---|---|---|---|---|---|---|
| stream | 20223.56 | 2527.94 | 8 | 55 | 6.73 | 4.07E-06 |

Supplementary table 1: **Median duration preference across streams.** Type III ANOVA on LME model estimates with Satterthwaite's method for degrees of freedom.

| | Sum.Sq | Mean.Sq | NumDF | DenDF | F.value | Pr(>F) |
|---|---|---|---|---|---|---|
| ROI | 70624.75 | 1121.03 | 63 | 756 | 2.98 | 2.07E-12 |

Supplementary table 2: **Median duration preference across ROIs.** Type III ANOVA on LME model estimates with Satterthwaite's method for degrees of freedom.

| contrast | estimate | SE | df | t.ratio | p.value |
|---|---|---|---|---|---|
| AI - IPS | 2.96 | 4.60 | 55 | 0.64 | 1.00E+00 |
| AI - IF | 7.91 | 5.40 | 55 | 1.47 | 1.00E+00 |
| AI - IP | 4.94 | 4.78 | 55 | 1.03 | 1.00E+00 |
| AI - LV | -11.71 | 4.71 | 55 | -2.49 | 5.75E-01 |
| AI - (mot-som) | 8.60 | 5.16 | 55 | 1.67 | 1.00E+00 |
| AI - PM | -0.48 | 4.88 | 55 | -0.10 | 1.00E+00 |
| AI - SMA | 4.26 | 4.60 | 55 | 0.93 | 1.00E+00 |
| AI - VV | -7.78 | 5.00 | 55 | -1.56 | 1.00E+00 |
| IPS - IF | 4.95 | 4.13 | 55 | 1.20 | 1.00E+00 |
| IPS - IP | 1.98 | 3.28 | 55 | 0.60 | 1.00E+00 |
| IPS - LV | -14.67 | 3.18 | 55 | -4.62 | 8.48E-04 |
| IPS - (mot-som) | 5.64 | 3.81 | 55 | 1.48 | 1.00E+00 |
| IPS - PM | -3.44 | 3.42 | 55 | -1.01 | 1.00E+00 |
| IPS - SMA | 1.30 | 3.01 | 55 | 0.43 | 1.00E+00 |
| IPS - VV | -10.74 | 3.59 | 55 | -3.00 | 1.48E-01 |
| IF - IP | -2.97 | 4.33 | 55 | -0.69 | 1.00E+00 |
| IF - LV | -19.62 | 4.25 | 55 | -4.62 | 8.45E-04 |
| IF - (mot-som) | 0.69 | 4.74 | 55 | 0.15 | 1.00E+00 |
| IF - PM | -8.39 | 4.43 | 55 | -1.90 | 1.00E+00 |
| IF - SMA | -3.65 | 4.13 | 55 | -0.88 | 1.00E+00 |
| IF - VV | -15.69 | 4.56 | 55 | -3.44 | 4.02E-02 |
| IP - LV | -16.65 | 3.43 | 55 | -4.85 | 3.79E-04 |
| IP - (mot-som) | 3.66 | 4.03 | 55 | 0.91 | 1.00E+00 |
| IP - PM | -5.42 | 3.66 | 55 | -1.48 | 1.00E+00 |
| IP - SMA | -0.68 | 3.28 | 55 | -0.21 | 1.00E+00 |
| IP - VV | -12.72 | 3.82 | 55 | -3.33 | 5.55E-02 |
| LV - (mot-som) | 20.31 | 3.94 | 55 | 5.15 | 1.28E-04 |
| LV - PM | 11.23 | 3.56 | 55 | 3.15 | 9.41E-02 |
| LV - SMA | 15.97 | 3.18 | 55 | 5.03 | 2.01E-04 |

| | | | | | |
|---|---|---|---|---|---|
| LV - VV | 3.93 | 3.72 | 55 | 1.06 | 1.00E+00 |
| (mot-som) - PM | -9.08 | 4.14 | 55 | -2.20 | 1.00E+00 |
| (mot-som) - SMA | -4.34 | 3.81 | 55 | -1.14 | 1.00E+00 |
| (mot-som) - VV | -16.38 | 4.28 | 55 | -3.83 | 1.20E-02 |
| PM - SMA | 4.74 | 3.42 | 55 | 1.39 | 1.00E+00 |
| PM - VV | -7.30 | 3.93 | 55 | -1.86 | 1.00E+00 |
| SMA - VV | -12.04 | 3.59 | 55 | -3.36 | 5.17E-02 |

Supplementary table 3: **Median duration preference across streams.** Two-sided t contrasts between streams (Bonferroni-corrected) with Kenward-Roger's method for degrees of freedom.

| contrast | estimate | SE | df | t.ratio | p.value |
|---|---|---|---|---|---|
| 1_hand - LO1 | -40.49 | 7.60 | 756 | -5.33 | 2.67E-04 |
| 1_hand - LO2 | -44.83 | 7.60 | 756 | -5.90 | 1.12E-05 |
| 1_hand - PIT | -35.29 | 7.60 | 756 | -4.64 | 8.22E-03 |
| 1_hand - V3B | -35.04 | 7.60 | 756 | -4.61 | 9.55E-03 |
| 1_hand - V3CD | -35.47 | 7.60 | 756 | -4.67 | 7.35E-03 |
| 1_hand - V4 | -33.17 | 7.60 | 756 | -4.36 | 2.95E-02 |
| 1_hand - V4t | -38.46 | 7.60 | 756 | -5.06 | 1.06E-03 |
| 24dd - LO2 | -34.66 | 7.60 | 756 | -4.56 | 1.21E-02 |
| 24dv - LO2 | -34.52 | 7.60 | 756 | -4.54 | 1.31E-02 |
| 3a_hand - LO1 | -35.48 | 7.60 | 756 | -4.67 | 7.30E-03 |
| 3a_hand - LO2 | -39.82 | 7.60 | 756 | -5.24 | 4.26E-04 |
| 3a_hand - V4t | -33.45 | 7.60 | 756 | -4.40 | 2.50E-02 |
| 4_hand - LO1 | -32.81 | 7.60 | 756 | -4.32 | 3.63E-02 |
| 4_hand - LO2 | -37.15 | 7.60 | 756 | -4.89 | 2.53E-03 |
| 7PL - LO1 | -34.69 | 7.60 | 756 | -4.56 | 1.19E-02 |
| 7PL - LO2 | -39.03 | 7.60 | 756 | -5.13 | 7.29E-04 |
| 7PL - V4t | -32.66 | 7.60 | 756 | -4.30 | 3.96E-02 |
| 8BM - LO2 | -35.42 | 7.60 | 756 | -4.66 | 7.55E-03 |
| IFJp - LO2 | -32.88 | 7.60 | 756 | -4.33 | 3.48E-02 |
| IFSp - LO2 | -35.99 | 7.60 | 756 | -4.73 | 5.28E-03 |
| IPS1 - LO2 | -33.81 | 7.60 | 756 | -4.45 | 2.02E-02 |
| LO1 - OP1 | 36.69 | 7.60 | 756 | 4.83 | 3.39E-03 |
| LO1 - VIP | 34.63 | 7.60 | 756 | 4.55 | 1.23E-02 |
| LO2 - OP1 | 41.03 | 7.60 | 756 | 5.40 | 1.83E-04 |
| LO2 - p9_46v | 33.58 | 7.60 | 756 | 4.42 | 2.32E-02 |
| LO2 - TE2p | 32.42 | 7.60 | 756 | 4.26 | 4.56E-02 |
| LO2 - VIP | 38.97 | 7.60 | 756 | 5.13 | 7.60E-04 |
| OP1 - V4t | -34.66 | 7.60 | 756 | -4.56 | 1.20E-02 |
| V4t - VIP | 32.60 | 7.60 | 756 | 4.29 | 4.11E-02 |

Supplementary table 4: **Median duration preference across ROIs.** Two-sided t contrasts between ROIs (Bonferroni-corrected) with Kenward-Roger's method for degrees of freedom. Only significant contrasts are reported.

| | DFn | DFd | F | p | p<.05 | ges |
|---|---|---|---|---|---|---|
| stream | 8 | 96 | 0.00 | 1.00E+00 | | 0.00 |
| category | 4 | 48 | 5.45 | 1.00E-03 | * | 0.08 |
| stream:category | 32 | 384 | 6.40 | 8.66E-21 | * | 0.30 |

Supplementary table 5: **Categorization of duration preferences across streams.** Type III two-way repeated-measures ANOVA.

| | W | p | p<.05 |
|---|---|---|---|
| category | 0.45 | 5.19E-01 | |

Supplementary table 6: **Categorization of duration preferences across streams.** Mauchly's sphericity test.

| | GGe | DF[GG] | p[GG] | p[GG]<.05 | HFe | DF[HF] | p[HF] | p[HF]<.05 |
|---|---|---|---|---|---|---|---|---|
| category | 0.72 | 2.86, 34.35 | 4.00E-03 | * | 0.96 | 3.85, 46.25 | 1.00E-03 | * |

Supplementary table 7: **Categorization of duration preferences across streams.** Deviation from sphericity correction. GGe = Greenhouse-Geisser $\epsilon$, HFe = Huynh-Feldt $\epsilon$.

| | DFn | DFd | F | p | p<.05 | ges |
|---|---|---|---|---|---|---|
| ROI | 63 | 756 | -0.00 | 1.00E+00 | | 0.00 |
| category | 4 | 48 | 4.38 | 4.00E-03 | * | 0.03 |
| ROI:category | 252 | 3024 | 3.15 | 2.11E-48 | * | 0.20 |

Supplementary table 8: **Categorization of duration preferences across ROIs.** Type III two-way repeated-measures ANOVA.

| | W | p | p<.05 |
|---|---|---|---|
| category | 0.42 | 4.69E-01 | |

Supplementary table 9: **Categorization of duration preferences across ROIs.** Mauchly's sphericity test.

| | GGe | DF[GG] | p[GG] | p[GG]<.05 | HFe | DF[HF] | p[HF] | p[HF]<.05 |
|---|---|---|---|---|---|---|---|---|
| category | 0.69 | 2.75, 33.06 | 1.20E-02 | * | 0.91 | 3.66, 43.89 | 6.00E-03 | * |

Supplementary table 10: **Categorization of duration preferences across ROIs.** Deviation from sphericity correction. GGe = Greenhouse-Geisser $\epsilon$, HFe = Huynh-Feldt $\epsilon$.

| contrast | stream | estimate | SE | df | t.ratio | p.value |
|---|---|---|---|---|---|---|
| short - (mid-short) | AI | -0.21 | 0.04 | 540 | -5.06 | 5.77E-06 |
| short - medium | AI | -0.39 | 0.04 | 540 | -9.67 | 1.72E-19 |
| short - (mid-long) | AI | -0.16 | 0.04 | 540 | -4.04 | 6.24E-04 |
| short - long | AI | -0.17 | 0.04 | 540 | -4.18 | 3.38E-04 |
| (mid-short) - medium | AI | -0.19 | 0.04 | 540 | -4.61 | 5.13E-05 |
| (mid-short) - (mid-long) | AI | 0.04 | 0.04 | 540 | 1.02 | 1.00E+00 |
| (mid-short) - long | AI | 0.04 | 0.04 | 540 | 0.88 | 1.00E+00 |
| medium - (mid-long) | AI | 0.23 | 0.04 | 540 | 5.63 | 2.90E-07 |
| medium - long | AI | 0.22 | 0.04 | 540 | 5.48 | 6.40E-07 |
| (mid-long) - long | AI | -0.01 | 0.04 | 540 | -0.15 | 1.00E+00 |
| short - (mid-short) | IPS | -0.04 | 0.04 | 540 | -0.89 | 1.00E+00 |
| short - medium | IPS | -0.01 | 0.04 | 540 | -0.13 | 1.00E+00 |
| short - (mid-long) | IPS | 0.01 | 0.04 | 540 | 0.24 | 1.00E+00 |
| short - long | IPS | -0.08 | 0.04 | 540 | -1.86 | 6.30E-01 |
| (mid-short) - medium | IPS | 0.03 | 0.04 | 540 | 0.76 | 1.00E+00 |
| (mid-short) - (mid-long) | IPS | 0.05 | 0.04 | 540 | 1.12 | 1.00E+00 |
| (mid-short) - long | IPS | -0.04 | 0.04 | 540 | -0.98 | 1.00E+00 |
| medium - (mid-long) | IPS | 0.01 | 0.04 | 540 | 0.37 | 1.00E+00 |
| medium - long | IPS | -0.07 | 0.04 | 540 | -1.73 | 8.35E-01 |
| (mid-long) - long | IPS | -0.09 | 0.04 | 540 | -2.10 | 3.60E-01 |
| short - (mid-short) | IF | -0.16 | 0.04 | 540 | -3.98 | 7.81E-04 |
| short - medium | IF | -0.22 | 0.04 | 540 | -5.49 | 6.29E-07 |
| short - (mid-long) | IF | -0.02 | 0.04 | 540 | -0.56 | 1.00E+00 |
| short - long | IF | -0.01 | 0.04 | 540 | -0.21 | 1.00E+00 |
| (mid-short) - medium | IF | -0.06 | 0.04 | 540 | -1.51 | 1.00E+00 |
| (mid-short) - (mid-long) | IF | 0.14 | 0.04 | 540 | 3.42 | 6.71E-03 |
| (mid-short) - long | IF | 0.15 | 0.04 | 540 | 3.77 | 1.81E-03 |
| medium - (mid-long) | IF | 0.20 | 0.04 | 540 | 4.93 | 1.11E-05 |
| medium - long | IF | 0.21 | 0.04 | 540 | 5.28 | 1.91E-06 |
| (mid-long) - long | IF | 0.01 | 0.04 | 540 | 0.35 | 1.00E+00 |
| short - (mid-short) | IP | 0.04 | 0.04 | 540 | 1.00 | 1.00E+00 |
| short - medium | IP | 0.06 | 0.04 | 540 | 1.40 | 1.00E+00 |
| short - (mid-long) | IP | 0.04 | 0.04 | 540 | 1.03 | 1.00E+00 |
| short - long | IP | 0.02 | 0.04 | 540 | 0.49 | 1.00E+00 |
| (mid-short) - medium | IP | 0.02 | 0.04 | 540 | 0.40 | 1.00E+00 |
| (mid-short) - (mid-long) | IP | 0.00 | 0.04 | 540 | 0.04 | 1.00E+00 |
| (mid-short) - long | IP | -0.02 | 0.04 | 540 | -0.50 | 1.00E+00 |
| medium - (mid-long) | IP | -0.01 | 0.04 | 540 | -0.37 | 1.00E+00 |
| medium - long | IP | -0.04 | 0.04 | 540 | -0.91 | 1.00E+00 |
| (mid-long) - long | IP | -0.02 | 0.04 | 540 | -0.54 | 1.00E+00 |
| short - (mid-short) | LV | -0.01 | 0.04 | 540 | -0.29 | 1.00E+00 |

| | | | | | | |
|---|---|---|---|---|---|---|
| short - medium | LV | -0.02 | 0.04 | 540 | -0.49 | 1.00E+00 |
| short - (mid-long) | LV | -0.05 | 0.04 | 540 | -1.24 | 1.00E+00 |
| short - long | LV | -0.22 | 0.04 | 540 | -5.31 | 1.63E-06 |
| (mid-short) - medium | LV | -0.01 | 0.04 | 540 | -0.20 | 1.00E+00 |
| (mid-short) - (mid-long) | LV | -0.04 | 0.04 | 540 | -0.95 | 1.00E+00 |
| (mid-short) - long | LV | -0.20 | 0.04 | 540 | -5.01 | 7.24E-06 |
| medium - (mid-long) | LV | -0.03 | 0.04 | 540 | -0.75 | 1.00E+00 |
| medium - long | LV | -0.20 | 0.04 | 540 | -4.81 | 1.93E-05 |
| (mid-long) - long | LV | -0.17 | 0.04 | 540 | -4.06 | 5.57E-04 |
| short - (mid-short) | mot-som | 0.09 | 0.04 | 540 | 2.17 | 3.07E-01 |
| short - medium | mot-som | 0.19 | 0.04 | 540 | 4.69 | 3.44E-05 |
| short - (mid-long) | mot-som | 0.17 | 0.04 | 540 | 4.24 | 2.67E-04 |
| short - long | mot-som | 0.04 | 0.04 | 540 | 0.87 | 1.00E+00 |
| (mid-short) - medium | mot-som | 0.10 | 0.04 | 540 | 2.53 | 1.18E-01 |
| (mid-short) - (mid-long) | mot-som | 0.08 | 0.04 | 540 | 2.07 | 3.89E-01 |
| (mid-short) - long | mot-som | -0.05 | 0.04 | 540 | -1.29 | 1.00E+00 |
| medium - (mid-long) | mot-som | -0.02 | 0.04 | 540 | -0.46 | 1.00E+00 |
| medium - long | mot-som | -0.16 | 0.04 | 540 | -3.82 | 1.49E-03 |
| (mid-long) - long | mot-som | -0.14 | 0.04 | 540 | -3.36 | 8.24E-03 |
| short - (mid-short) | PM | -0.05 | 0.04 | 540 | -1.24 | 1.00E+00 |
| short - medium | PM | -0.10 | 0.04 | 540 | -2.51 | 1.24E-01 |
| short - (mid-long) | PM | -0.05 | 0.04 | 540 | -1.13 | 1.00E+00 |
| short - long | PM | -0.10 | 0.04 | 540 | -2.53 | 1.16E-01 |
| (mid-short) - medium | PM | -0.05 | 0.04 | 540 | -1.27 | 1.00E+00 |
| (mid-short) - (mid-long) | PM | 0.00 | 0.04 | 540 | 0.11 | 1.00E+00 |
| (mid-short) - long | PM | -0.05 | 0.04 | 540 | -1.29 | 1.00E+00 |
| medium - (mid-long) | PM | 0.06 | 0.04 | 540 | 1.38 | 1.00E+00 |
| medium - long | PM | -0.00 | 0.04 | 540 | -0.02 | 1.00E+00 |
| (mid-long) - long | PM | -0.06 | 0.04 | 540 | -1.41 | 1.00E+00 |
| short - (mid-short) | SMA | -0.04 | 0.04 | 540 | -0.99 | 1.00E+00 |
| short - medium | SMA | -0.11 | 0.04 | 540 | -2.71 | 6.85E-02 |
| short - (mid-long) | SMA | 0.01 | 0.04 | 540 | 0.20 | 1.00E+00 |
| short - long | SMA | -0.01 | 0.04 | 540 | -0.31 | 1.00E+00 |
| (mid-short) - medium | SMA | -0.07 | 0.04 | 540 | -1.73 | 8.43E-01 |
| (mid-short) - (mid-long) | SMA | 0.05 | 0.04 | 540 | 1.19 | 1.00E+00 |
| (mid-short) - long | SMA | 0.03 | 0.04 | 540 | 0.68 | 1.00E+00 |
| medium - (mid-long) | SMA | 0.12 | 0.04 | 540 | 2.91 | 3.71E-02 |
| medium - long | SMA | 0.10 | 0.04 | 540 | 2.41 | 1.63E-01 |
| (mid-long) - long | SMA | -0.02 | 0.04 | 540 | -0.51 | 1.00E+00 |
| short - (mid-short) | VV | -0.01 | 0.04 | 540 | -0.36 | 1.00E+00 |
| short - medium | VV | -0.02 | 0.04 | 540 | -0.44 | 1.00E+00 |
| short - (mid-long) | VV | -0.06 | 0.04 | 540 | -1.38 | 1.00E+00 |

| | | | | | | |
|---|---|---|---|---|---|---|
| short - long | VV | -0.16 | 0.04 | 540 | -3.83 | 1.45E-03 |
| (mid-short) - medium | VV | -0.00 | 0.04 | 540 | -0.08 | 1.00E+00 |
| (mid-short) - (mid-long) | VV | -0.04 | 0.04 | 540 | -1.03 | 1.00E+00 |
| (mid-short) - long | VV | -0.14 | 0.04 | 540 | -3.47 | 5.58E-03 |
| medium - (mid-long) | VV | -0.04 | 0.04 | 540 | -0.95 | 1.00E+00 |
| medium - long | VV | -0.14 | 0.04 | 540 | -3.39 | 7.46E-03 |
| (mid-long) - long | VV | -0.10 | 0.04 | 540 | -2.44 | 1.49E-01 |

Supplementary table 11: **Categorization of duration preferences across streams.** Two-sided t contrasts between categories for each stream (Bonferroni-corrected). Contrast estimates are expressed as difference of square-root scaled values.

| contrast | ROI | estimate | SE | df | t.ratio | p.value |
|---|---|---|---|---|---|---|
| short - medium | 1_hand | 0.19 | 0.07 | 3840 | 2.95 | 3.23E-02 |
| short - (mid-long) | 1_hand | 0.20 | 0.07 | 3840 | 3.10 | 1.95E-02 |
| short - (mid-long) | 24dd | 0.19 | 0.07 | 3840 | 2.87 | 4.16E-02 |
| short - medium | 3a_hand | 0.29 | 0.07 | 3840 | 4.41 | 1.07E-04 |
| short - (mid-long) | 3a_hand | 0.28 | 0.07 | 3840 | 4.23 | 2.40E-04 |
| medium - long | 3a_hand | -0.20 | 0.07 | 3840 | -3.10 | 1.93E-02 |
| (mid-long) - long | 3a_hand | -0.19 | 0.07 | 3840 | -2.92 | 3.47E-02 |
| short - medium | 3b_hand | 0.19 | 0.07 | 3840 | 2.85 | 4.33E-02 |
| short - (mid-long) | 3b_hand | 0.20 | 0.07 | 3840 | 3.06 | 2.24E-02 |
| medium - long | 3b_hand | -0.19 | 0.07 | 3840 | -2.81 | 4.94E-02 |
| (mid-long) - long | 3b_hand | -0.20 | 0.07 | 3840 | -3.02 | 2.57E-02 |
| short - medium | 4_hand | 0.24 | 0.07 | 3840 | 3.67 | 2.48E-03 |
| short - (mid-long) | 4_hand | 0.21 | 0.07 | 3840 | 3.24 | 1.21E-02 |
| short - long | 55b | -0.25 | 0.07 | 3840 | -3.82 | 1.35E-03 |
| (mid-long) - long | 55b | -0.21 | 0.07 | 3840 | -3.16 | 1.57E-02 |
| (mid-short) - long | 6v | -0.20 | 0.07 | 3840 | -3.10 | 1.93E-02 |
| short - medium | 8BM | -0.26 | 0.07 | 3840 | -3.89 | 1.02E-03 |
| (mid-short) - long | 8BM | 0.22 | 0.07 | 3840 | 3.27 | 1.07E-02 |
| medium - (mid-long) | 8BM | 0.24 | 0.07 | 3840 | 3.62 | 3.01E-03 |
| medium - long | 8BM | 0.38 | 0.07 | 3840 | 5.68 | 1.42E-07 |
| short - medium | a24pr | -0.25 | 0.07 | 3840 | -3.81 | 1.42E-03 |
| short - medium | a32pr | -0.33 | 0.07 | 3840 | -4.99 | 6.23E-06 |
| (mid-short) - medium | a32pr | -0.22 | 0.07 | 3840 | -3.33 | 8.62E-03 |
| medium - long | a32pr | 0.27 | 0.07 | 3840 | 4.16 | 3.31E-04 |
| short - (mid-short) | AVI | -0.32 | 0.07 | 3840 | -4.82 | 1.50E-05 |
| short - medium | AVI | -0.43 | 0.07 | 3840 | -6.53 | 7.39E-10 |
| (mid-short) - (mid-long) | AVI | 0.20 | 0.07 | 3840 | 3.07 | 2.15E-02 |
| (mid-short) - long | AVI | 0.25 | 0.07 | 3840 | 3.81 | 1.38E-03 |
| medium - (mid-long) | AVI | 0.32 | 0.07 | 3840 | 4.78 | 1.79E-05 |
| medium - long | AVI | 0.37 | 0.07 | 3840 | 5.53 | 3.46E-07 |
| short - medium | FOP4 | -0.31 | 0.07 | 3840 | -4.76 | 1.99E-05 |
| (mid-short) - medium | FOP4 | -0.21 | 0.07 | 3840 | -3.16 | 1.60E-02 |
| short - (mid-short) | FOP5 | -0.20 | 0.07 | 3840 | -2.99 | 2.84E-02 |
| short - medium | FOP5 | -0.44 | 0.07 | 3840 | -6.67 | 2.93E-10 |
| short - (mid-long) | FOP5 | -0.21 | 0.07 | 3840 | -3.23 | 1.24E-02 |
| short - long | FOP5 | -0.25 | 0.07 | 3840 | -3.82 | 1.36E-03 |
| (mid-short) - medium | FOP5 | -0.24 | 0.07 | 3840 | -3.68 | 2.33E-03 |
| medium - (mid-long) | FOP5 | 0.23 | 0.07 | 3840 | 3.44 | 5.92E-03 |
| medium - long | FOP5 | 0.19 | 0.07 | 3840 | 2.85 | 4.39E-02 |
| short - (mid-short) | IFJp | -0.24 | 0.07 | 3840 | -3.71 | 2.13E-03 |
| short - medium | IFJp | -0.29 | 0.07 | 3840 | -4.39 | 1.14E-04 |

| | | | | | | |
|---|---|---|---|---|---|---|
| (mid-short) - (mid-long) | IFJp | 0.21 | 0.07 | 3840 | 3.25 | 1.15E-02 |
| (mid-short) - long | IFJp | 0.21 | 0.07 | 3840 | 3.25 | 1.15E-02 |
| medium - (mid-long) | IFJp | 0.26 | 0.07 | 3840 | 3.94 | 8.26E-04 |
| medium - long | IFJp | 0.26 | 0.07 | 3840 | 3.94 | 8.27E-04 |
| (mid-short) - (mid-long) | IFSp | 0.20 | 0.07 | 3840 | 3.01 | 2.65E-02 |
| (mid-short) - long | IFSp | 0.23 | 0.07 | 3840 | 3.55 | 3.90E-03 |
| short - long | LIPv | -0.24 | 0.07 | 3840 | -3.57 | 3.60E-03 |
| short - long | LO1 | -0.32 | 0.07 | 3840 | -4.90 | 9.82E-06 |
| (mid-short) - long | LO1 | -0.41 | 0.07 | 3840 | -6.14 | 8.88E-09 |
| medium - long | LO1 | -0.42 | 0.07 | 3840 | -6.32 | 2.85E-09 |
| (mid-long) - long | LO1 | -0.33 | 0.07 | 3840 | -5.03 | 5.17E-06 |
| short - long | LO2 | -0.34 | 0.07 | 3840 | -5.17 | 2.47E-06 |
| (mid-short) - long | LO2 | -0.38 | 0.07 | 3840 | -5.80 | 7.12E-08 |
| medium - long | LO2 | -0.39 | 0.07 | 3840 | -5.83 | 5.84E-08 |
| (mid-long) - long | LO2 | -0.32 | 0.07 | 3840 | -4.86 | 1.20E-05 |
| short - long | MST | -0.20 | 0.07 | 3840 | -3.05 | 2.31E-02 |
| short - medium | OP1 | 0.21 | 0.07 | 3840 | 3.14 | 1.68E-02 |
| short - (mid-long) | OP1 | 0.19 | 0.07 | 3840 | 2.94 | 3.30E-02 |
| short - medium | p32pr | -0.37 | 0.07 | 3840 | -5.67 | 1.56E-07 |
| short - (mid-long) | p32pr | -0.22 | 0.07 | 3840 | -3.36 | 7.90E-03 |
| (mid-short) - medium | p32pr | -0.21 | 0.07 | 3840 | -3.17 | 1.55E-02 |
| medium - long | p32pr | 0.26 | 0.07 | 3840 | 3.87 | 1.08E-03 |
| short - (mid-short) | p9_46v | -0.20 | 0.07 | 3840 | -2.96 | 3.13E-02 |
| short - medium | p9_46v | -0.31 | 0.07 | 3840 | -4.73 | 2.33E-05 |
| (mid-short) - long | p9_46v | 0.20 | 0.07 | 3840 | 3.02 | 2.58E-02 |
| medium - (mid-long) | p9_46v | 0.27 | 0.07 | 3840 | 4.07 | 4.81E-04 |
| medium - long | p9_46v | 0.32 | 0.07 | 3840 | 4.79 | 1.74E-05 |
| short - medium | PEF | -0.26 | 0.07 | 3840 | -4.00 | 6.44E-04 |
| short - long | PIT | -0.27 | 0.07 | 3840 | -4.04 | 5.45E-04 |
| (mid-short) - long | PIT | -0.25 | 0.07 | 3840 | -3.77 | 1.68E-03 |
| medium - long | PIT | -0.28 | 0.07 | 3840 | -4.27 | 2.00E-04 |
| short - medium | SCEF | -0.29 | 0.07 | 3840 | -4.42 | 1.00E-04 |
| (mid-short) - medium | SCEF | -0.21 | 0.07 | 3840 | -3.13 | 1.77E-02 |
| medium - (mid-long) | SCEF | 0.20 | 0.07 | 3840 | 3.02 | 2.53E-02 |
| medium - long | SFL | 0.19 | 0.07 | 3840 | 2.93 | 3.43E-02 |
| short - long | V3B | -0.29 | 0.07 | 3840 | -4.43 | 9.91E-05 |
| (mid-short) - long | V3B | -0.28 | 0.07 | 3840 | -4.20 | 2.74E-04 |
| medium - long | V3B | -0.40 | 0.07 | 3840 | -6.04 | 1.66E-08 |
| (mid-long) - long | V3B | -0.29 | 0.07 | 3840 | -4.46 | 8.51E-05 |
| short - long | V3CD | -0.25 | 0.07 | 3840 | -3.76 | 1.73E-03 |
| (mid-short) - long | V3CD | -0.24 | 0.07 | 3840 | -3.66 | 2.59E-03 |
| medium - long | V3CD | -0.27 | 0.07 | 3840 | -4.06 | 5.09E-04 |

| | | | | | | |
|---|---|---|---|---|---|---|
| (mid-long) - long | V3CD | -0.26 | 0.07 | 3840 | -3.90 | 9.60E-04 |
| short - long | V4 | -0.23 | 0.07 | 3840 | -3.43 | 6.06E-03 |
| medium - long | V4 | -0.20 | 0.07 | 3840 | -3.00 | 2.74E-02 |
| short - long | V4t | -0.30 | 0.07 | 3840 | -4.58 | 4.72E-05 |
| medium - long | V4t | -0.20 | 0.07 | 3840 | -2.99 | 2.82E-02 |
| (mid-short) - (mid-long) | V8 | -0.19 | 0.07 | 3840 | -2.93 | 3.42E-02 |
| short - long | VVC | -0.22 | 0.07 | 3840 | -3.31 | 9.57E-03 |

Supplementary table 12: **Categorization of duration preferences across ROIs.** Two-sided t contrasts between categories for each ROI (Bonferroni-corrected). Contrast estimates are expressed as difference of square-root scaled values. Only significant contrasts are reported.

| | Sum.Sq | Mean.Sq | NumDF | DenDF | F.value | Pr(>F) |
|---|---|---|---|---|---|---|
| stream | 2.39 | 0.30 | 8 | 55 | 6.47 | 6.52E-06 |

Supplementary table 13: **Difference between fractions of vertices showing LL and HH types of spatial association across streams.** Type III ANOVA on LME model estimates with Satterthwaite's method for degrees of freedom.

| | Sum.Sq | Mean.Sq | NumDF | DenDF | F.value | Pr(>F) |
|---|---|---|---|---|---|---|
| ROI | 8.15 | 0.13 | 63 | 756 | 2.81 | 4.19E-11 |

Supplementary table 14: **Difference between fractions of vertices showing LL and HH types of spatial association across ROIs.** Type III ANOVA on LME model estimates with Satterthwaite's method for degrees of freedom.

| | DFn | DFd | F | p | p<.05 | ges |
|---|---|---|---|---|---|---|
| stream | 8 | 96 | 1.20 | 3.08E-01 | | 0.01 |
| type | 1 | 12 | 1.92 | 1.91E-01 | | 0.02 |
| stream:type | 8 | 96 | 3.87 | 5.54E-04 | * | 0.19 |

Supplementary table 15: **Fractions of vertices showing LL and HH types of spatial association across streams.** Type III two-way repeated-measures ANOVA.

| | W | p | p<.05 |
|---|---|---|---|
| stream | 0.00 | 7.00E-03 | * |
| stream:type | 0.00 | 2.00E-03 | * |

Supplementary table 16: **Fractions of vertices showing LL and HH types of spatial association across streams.** Mauchly's sphericity test.

| | GGe | DF[GG] | p[GG] | p[GG]<.05 | HFe | DF[HF] | p[HF] | p[HF]<.05 |
|---|---|---|---|---|---|---|---|---|
| stream | 0.44 | 3.55, 42.61 | 3.24E-01 | | 0.65 | 5.23, 62.72 | 3.20E-01 | |
| stream:type | 0.46 | 3.71, 44.46 | 1.00E-02 | * | 0.70 | 5.57, 66.79 | 3.00E-03 | * |

Supplementary table 17: **Fractions of vertices showing LL and HH types of spatial association across streams.** Deviation from sphericity correction. GGe = Greenhouse-Geisser $\epsilon$, HFe = Huynh-Feldt $\epsilon$.

| | DFn | DFd | F | p | p<.05 | ges |
|---|---|---|---|---|---|---|
| ROI | 63 | 756 | 1.60 | 3.00E-03 | * | 0.02 |
| type | 1 | 12 | 3.88 | 7.30E-02 | | 0.01 |
| ROI:type | 63 | 756 | 2.81 | 4.19E-11 | * | 0.16 |

Supplementary table 18: **Fractions of vertices showing LL and HH types of spatial association across ROIs.** Type III two-way repeated-measures ANOVA. Mauchly's test of sphericity could not be evaluated due to the large number of within-subject levels.

| contrast | areas | estimate | SE | df | t.ratio | p.value |
|---|---|---|---|---|---|---|
| HH - LL | AI | 0.05 | 0.04 | 216 | 1.28 | 2.03E-01 |
| HH - LL | IPS | 0.01 | 0.04 | 216 | 0.27 | 7.91E-01 |
| HH - LL | IF | -0.10 | 0.04 | 216 | -2.67 | 8.09E-03 |
| HH - LL | IP | -0.00 | 0.04 | 216 | -0.08 | 9.37E-01 |
| HH - LL | LV | 0.18 | 0.04 | 216 | 4.98 | 1.28E-06 |
| HH - LL | mot-som | -0.08 | 0.04 | 216 | -2.16 | 3.22E-02 |
| HH - LL | PM | 0.06 | 0.04 | 216 | 1.58 | 1.16E-01 |
| HH - LL | SMA | -0.01 | 0.04 | 216 | -0.36 | 7.18E-01 |
| HH - LL | VV | 0.14 | 0.04 | 216 | 4.03 | 7.72E-05 |

Supplementary table 19: **Fractions of vertices showing LL and HH types of spatial association across streams.** Two-sided t contrasts between HH and LL spatial associations for each stream (Bonferroni-corrected).

| contrast | ROI | estimate | SE | df | t.ratio | p.value |
|---|---|---|---|---|---|---|
| HH - LL | 1_hand | -0.18 | 0.06 | 1536 | -2.93 | 3.48E-03 |
| HH - LL | 24dd | -0.13 | 0.06 | 1536 | -2.22 | 2.62E-02 |
| HH - LL | 24dv | -0.12 | 0.06 | 1536 | -1.94 | 5.30E-02 |
| HH - LL | 3a_hand | -0.14 | 0.06 | 1536 | -2.28 | 2.25E-02 |
| HH - LL | 3b_hand | -0.09 | 0.06 | 1536 | -1.44 | 1.50E-01 |
| HH - LL | 4_face | 0.13 | 0.06 | 1536 | 2.20 | 2.79E-02 |
| HH - LL | 4_hand | -0.10 | 0.06 | 1536 | -1.65 | 9.94E-02 |
| HH - LL | 44 | 0.05 | 0.06 | 1536 | 0.85 | 3.96E-01 |
| HH - LL | 55b | 0.12 | 0.06 | 1536 | 2.03 | 4.28E-02 |
| HH - LL | 6a | -0.02 | 0.06 | 1536 | -0.41 | 6.81E-01 |
| HH - LL | 6d | -0.01 | 0.06 | 1536 | -0.23 | 8.18E-01 |
| HH - LL | 6ma | 0.08 | 0.06 | 1536 | 1.27 | 2.05E-01 |
| HH - LL | 6mp | 0.06 | 0.06 | 1536 | 0.97 | 3.34E-01 |
| HH - LL | 6r | 0.06 | 0.06 | 1536 | 1.04 | 2.98E-01 |
| HH - LL | 6v | 0.16 | 0.06 | 1536 | 2.69 | 7.32E-03 |
| HH - LL | 7PL | -0.14 | 0.06 | 1536 | -2.39 | 1.69E-02 |
| HH - LL | 8BM | -0.19 | 0.06 | 1536 | -3.08 | 2.07E-03 |
| HH - LL | a24pr | 0.12 | 0.06 | 1536 | 2.01 | 4.46E-02 |
| HH - LL | a32pr | 0.04 | 0.06 | 1536 | 0.75 | 4.56E-01 |
| HH - LL | AIP | -0.00 | 0.06 | 1536 | -0.05 | 9.61E-01 |
| HH - LL | AVI | -0.12 | 0.06 | 1536 | -1.94 | 5.25E-02 |
| HH - LL | FEF | 0.10 | 0.06 | 1536 | 1.73 | 8.33E-02 |
| HH - LL | FFC | 0.13 | 0.06 | 1536 | 2.12 | 3.45E-02 |
| HH - LL | FOP4 | 0.16 | 0.06 | 1536 | 2.60 | 9.32E-03 |
| HH - LL | FOP5 | 0.11 | 0.06 | 1536 | 1.91 | 5.68E-02 |
| HH - LL | FST | 0.11 | 0.06 | 1536 | 1.88 | 6.10E-02 |
| HH - LL | IFJp | -0.14 | 0.06 | 1536 | -2.32 | 2.05E-02 |
| HH - LL | IFSp | -0.22 | 0.06 | 1536 | -3.66 | 2.62E-04 |
| HH - LL | IP0 | -0.03 | 0.06 | 1536 | -0.43 | 6.65E-01 |
| HH - LL | IP2 | -0.04 | 0.06 | 1536 | -0.59 | 5.58E-01 |

| | | | | | | |
|---|---|---|---|---|---|---|
| HH - LL | IPS1 | -0.12 | 0.06 | 1536 | -1.92 | 5.57E-02 |
| HH - LL | LIPd | 0.01 | 0.06 | 1536 | 0.08 | 9.34E-01 |
| HH - LL | LIPv | 0.15 | 0.06 | 1536 | 2.43 | 1.53E-02 |
| HH - LL | LO1 | 0.32 | 0.06 | 1536 | 5.31 | 1.23E-07 |
| HH - LL | LO2 | 0.33 | 0.06 | 1536 | 5.42 | 7.06E-08 |
| HH - LL | LO3 | 0.11 | 0.06 | 1536 | 1.87 | 6.14E-02 |
| HH - LL | MIP | 0.06 | 0.06 | 1536 | 1.04 | 2.99E-01 |
| HH - LL | MST | 0.20 | 0.06 | 1536 | 3.36 | 7.90E-04 |
| HH - LL | MT | 0.14 | 0.06 | 1536 | 2.39 | 1.69E-02 |
| HH - LL | OP1 | -0.13 | 0.06 | 1536 | -2.18 | 2.93E-02 |
| HH - LL | p24pr | -0.02 | 0.06 | 1536 | -0.25 | 8.01E-01 |
| HH - LL | p32pr | 0.15 | 0.06 | 1536 | 2.49 | 1.28E-02 |
| HH - LL | p9_46v | -0.12 | 0.06 | 1536 | -1.96 | 5.01E-02 |
| HH - LL | PEF | 0.02 | 0.06 | 1536 | 0.36 | 7.22E-01 |
| HH - LL | PFop | -0.04 | 0.06 | 1536 | -0.60 | 5.51E-01 |
| HH - LL | PFt | -0.03 | 0.06 | 1536 | -0.49 | 6.23E-01 |
| HH - LL | PGp | -0.03 | 0.06 | 1536 | -0.48 | 6.29E-01 |
| HH - LL | PH | 0.14 | 0.06 | 1536 | 2.26 | 2.40E-02 |
| HH - LL | PIT | 0.28 | 0.06 | 1536 | 4.72 | 2.51E-06 |
| HH - LL | SCEF | 0.09 | 0.06 | 1536 | 1.53 | 1.27E-01 |
| HH - LL | SFL | -0.05 | 0.06 | 1536 | -0.81 | 4.17E-01 |
| HH - LL | STV | 0.00 | 0.06 | 1536 | 0.03 | 9.80E-01 |
| HH - LL | TE2p | -0.05 | 0.06 | 1536 | -0.81 | 4.20E-01 |
| HH - LL | TPOJ1 | 0.03 | 0.06 | 1536 | 0.52 | 6.03E-01 |
| HH - LL | TPOJ2 | 0.04 | 0.06 | 1536 | 0.70 | 4.82E-01 |
| HH - LL | TPOJ3 | 0.02 | 0.06 | 1536 | 0.36 | 7.17E-01 |
| HH - LL | V3B | 0.17 | 0.06 | 1536 | 2.76 | 5.82E-03 |
| HH - LL | V3CD | 0.18 | 0.06 | 1536 | 2.94 | 3.31E-03 |
| HH - LL | V4 | 0.17 | 0.06 | 1536 | 2.81 | 5.03E-03 |
| HH - LL | V4t | 0.27 | 0.06 | 1536 | 4.42 | 1.04E-05 |
| HH - LL | V7 | -0.02 | 0.06 | 1536 | -0.33 | 7.42E-01 |
| HH - LL | V8 | 0.24 | 0.06 | 1536 | 3.99 | 6.98E-05 |
| HH - LL | VIP | -0.15 | 0.06 | 1536 | -2.54 | 1.11E-02 |
| HH - LL | VVC | 0.21 | 0.06 | 1536 | 3.44 | 5.90E-04 |

Supplementary table 20: **Fractions of vertices showing LL and HH types of spatial association across ROIs.** Two-sided t contrasts between HH and LL spatial associations for each ROI (Bonferroni-corrected).

| | Sum.Sq | Mean.Sq | NumDF | DenDF | F.value | Pr(>F) |
|---|---|---|---|---|---|---|
| stream | 0.13 | 0.02 | 8 | 96 | 10.26 | 2.90E-10 |

Supplementary table 21: **Nuggets across streams.** Type III ANOVA on LME model estimates with Satterthwaite's method for degrees of freedom.

| | Sum.Sq | Mean.Sq | NumDF | DenDF | F.value | Pr(>F) |
|---|---|---|---|---|---|---|
| stream | 0.12 | 0.02 | 8 | 96 | 15.76 | 1.31E-14 |

Supplementary table 22: **Ranges across streams.** Type III ANOVA on LME model estimates with Satterthwaite's method for degrees of freedom.

| contrast | estimate | SE | df | t.ratio | p.value |
|---|---|---|---|---|---|
| VV - AI | 0.08 | 0.02 | 96 | 5.00 | 9.13E-05 |
| VV - IPS | 0.08 | 0.02 | 96 | 5.33 | 2.35E-05 |
| VV - IF | 0.05 | 0.02 | 96 | 2.91 | 1.61E-01 |
| VV - IP | 0.04 | 0.02 | 96 | 2.42 | 6.25E-01 |
| VV - LV | 0.01 | 0.02 | 96 | 0.94 | 1.00E+00 |
| VV - (mot-som) | 0.11 | 0.02 | 96 | 7.20 | 4.95E-09 |
| VV - PM | 0.03 | 0.02 | 96 | 2.20 | 1.00E+00 |
| VV - SMA | 0.06 | 0.02 | 96 | 3.80 | 9.18E-03 |
| AI - IPS | 0.01 | 0.02 | 96 | 0.32 | 1.00E+00 |
| AI - IF | -0.03 | 0.02 | 96 | -2.09 | 1.00E+00 |
| AI - IP | -0.04 | 0.02 | 96 | -2.58 | 4.07E-01 |
| AI - LV | -0.06 | 0.02 | 96 | -4.06 | 3.55E-03 |
| AI - (mot-som) | 0.03 | 0.02 | 96 | 2.19 | 1.00E+00 |
| AI - PM | -0.04 | 0.02 | 96 | -2.80 | 2.21E-01 |
| AI - SMA | -0.02 | 0.02 | 96 | -1.21 | 1.00E+00 |
| IPS - IF | -0.04 | 0.02 | 96 | -2.42 | 6.33E-01 |
| IPS - IP | -0.05 | 0.02 | 96 | -2.91 | 1.63E-01 |
| IPS - LV | -0.07 | 0.02 | 96 | -4.39 | 1.05E-03 |
| IPS - (mot-som) | 0.03 | 0.02 | 96 | 1.87 | 1.00E+00 |
| IPS - PM | -0.05 | 0.02 | 96 | -3.13 | 8.44E-02 |
| IPS - SMA | -0.02 | 0.02 | 96 | -1.53 | 1.00E+00 |
| IF - IP | -0.01 | 0.02 | 96 | -0.49 | 1.00E+00 |
| IF - LV | -0.03 | 0.02 | 96 | -1.97 | 1.00E+00 |
| IF - (mot-som) | 0.07 | 0.02 | 96 | 4.28 | 1.58E-03 |
| IF - PM | -0.01 | 0.02 | 96 | -0.71 | 1.00E+00 |
| IF - SMA | 0.01 | 0.02 | 96 | 0.89 | 1.00E+00 |
| IP - LV | -0.02 | 0.02 | 96 | -1.48 | 1.00E+00 |
| IP - (mot-som) | 0.07 | 0.02 | 96 | 4.77 | 2.33E-04 |
| IP - PM | -0.00 | 0.02 | 96 | -0.22 | 1.00E+00 |
| IP - SMA | 0.02 | 0.02 | 96 | 1.38 | 1.00E+00 |
| LV - (mot-som) | 0.10 | 0.02 | 96 | 6.26 | 3.96E-07 |
| LV - PM | 0.02 | 0.02 | 96 | 1.26 | 1.00E+00 |
| LV - SMA | 0.04 | 0.02 | 96 | 2.86 | 1.88E-01 |
| (mot-som) - PM | -0.08 | 0.02 | 96 | -4.99 | 9.56E-05 |
| (mot-som) - SMA | -0.05 | 0.02 | 96 | -3.40 | 3.58E-02 |
| PM - SMA | 0.03 | 0.02 | 96 | 1.60 | 1.00E+00 |

Supplementary table 23: **Nuggets across streams.** Two-sided t contrasts between streams (Bonferroni-corrected) with Kenward-Roger's method for degrees of freedom.

| contrast | estimate | SE | df | t.ratio | p.value |
|---|---|---|---|---|---|
| VV - AI | -0.10 | 0.01 | 96 | -7.84 | 2.22E-10 |
| VV - IPS | -0.08 | 0.01 | 96 | -6.89 | 2.11E-08 |
| VV - IF | -0.10 | 0.01 | 96 | -8.31 | 2.29E-11 |
| VV - IP | -0.08 | 0.01 | 96 | -6.32 | 3.01E-07 |
| VV - LV | -0.07 | 0.01 | 96 | -5.64 | 6.18E-06 |
| VV - (mot-som) | -0.02 | 0.01 | 96 | -1.50 | 1.00E+00 |
| VV - PM | -0.07 | 0.01 | 96 | -5.76 | 3.68E-06 |
| VV - SMA | -0.05 | 0.01 | 96 | -4.14 | 2.70E-03 |
| AI - IPS | 0.01 | 0.01 | 96 | 0.95 | 1.00E+00 |
| AI - IF | -0.01 | 0.01 | 96 | -0.47 | 1.00E+00 |
| AI - IP | 0.02 | 0.01 | 96 | 1.52 | 1.00E+00 |
| AI - LV | 0.03 | 0.01 | 96 | 2.20 | 1.00E+00 |
| AI - (mot-som) | 0.08 | 0.01 | 96 | 6.34 | 2.71E-07 |
| AI - PM | 0.03 | 0.01 | 96 | 2.08 | 1.00E+00 |
| AI - SMA | 0.05 | 0.01 | 96 | 3.70 | 1.29E-02 |
| IPS - IF | -0.02 | 0.01 | 96 | -1.42 | 1.00E+00 |
| IPS - IP | 0.01 | 0.01 | 96 | 0.57 | 1.00E+00 |
| IPS - LV | 0.02 | 0.01 | 96 | 1.25 | 1.00E+00 |
| IPS - (mot-som) | 0.07 | 0.01 | 96 | 5.39 | 1.83E-05 |
| IPS - PM | 0.01 | 0.01 | 96 | 1.13 | 1.00E+00 |
| IPS - SMA | 0.03 | 0.01 | 96 | 2.75 | 2.56E-01 |
| IF - IP | 0.02 | 0.01 | 96 | 1.99 | 1.00E+00 |
| IF - LV | 0.03 | 0.01 | 96 | 2.67 | 3.24E-01 |
| IF - (mot-som) | 0.08 | 0.01 | 96 | 6.80 | 3.14E-08 |
| IF - PM | 0.03 | 0.01 | 96 | 2.55 | 4.47E-01 |
| IF - SMA | 0.05 | 0.01 | 96 | 4.17 | 2.43E-03 |
| IP - LV | 0.01 | 0.01 | 96 | 0.68 | 1.00E+00 |
| IP - (mot-som) | 0.06 | 0.01 | 96 | 4.81 | 1.98E-04 |
| IP - PM | 0.01 | 0.01 | 96 | 0.56 | 1.00E+00 |
| IP - SMA | 0.03 | 0.01 | 96 | 2.18 | 1.00E+00 |
| LV - (mot-som) | 0.05 | 0.01 | 96 | 4.14 | 2.71E-03 |
| LV - PM | -0.00 | 0.01 | 96 | -0.12 | 1.00E+00 |
| LV - SMA | 0.02 | 0.01 | 96 | 1.50 | 1.00E+00 |
| (mot-som) - PM | -0.05 | 0.01 | 96 | -4.26 | 1.75E-03 |
| (mot-som) - SMA | -0.03 | 0.01 | 96 | -2.64 | 3.52E-01 |
| PM - SMA | 0.02 | 0.01 | 96 | 1.62 | 1.00E+00 |

Supplementary table 24: **Ranges across streams.** Two-sided t contrasts between streams (Bonferroni-corrected) with Kenward-Roger's method for degrees of freedom. Contrast estimates are expressed as difference of square-root scaled values.

| | Sum.Sq | Mean.Sq | NumDF | DenDF | F.value | Pr(>F) |
|---|---|---|---|---|---|---|
| ROI | 2.22 | 0.04 | 63 | 756 | 4.40 | 2.55E-23 |

Supplementary table 25: **Nuggets across ROIs.** Type III ANOVA on LME model estimates with Satterthwaite's method for degrees of freedom.

| | Sum.Sq | Mean.Sq | NumDF | DenDF | F.value | Pr(>F) |
|---|---|---|---|---|---|---|
| ROI | 31.34 | 0.50 | 63 | 756 | 3.26 | 1.60E-14 |

Supplementary table 26: **Ranges across ROIs.** Type III ANOVA on LME model estimates with Satterthwaite's method for degrees of freedom.

| contrast | estimate | SE | df | t.ratio | p.value |
|---|---|---|---|---|---|
| 3a_hand - 6v | -0.20 | 0.04 | 756 | -5.72 | 3.07E-05 |
| 3a_hand - a24pr | -0.18 | 0.04 | 756 | -5.27 | 3.68E-04 |
| 3a_hand - FFC | -0.16 | 0.04 | 756 | -4.53 | 1.36E-02 |
| 3a_hand - LO2 | -0.20 | 0.04 | 756 | -5.61 | 5.73E-05 |
| 3a_hand - OP1 | -0.15 | 0.04 | 756 | -4.28 | 4.28E-02 |
| 3a_hand - p24pr | -0.18 | 0.04 | 756 | -5.02 | 1.32E-03 |
| 3a_hand - PIT | -0.20 | 0.04 | 756 | -5.78 | 2.21E-05 |
| 3a_hand - V3CD | -0.16 | 0.04 | 756 | -4.42 | 2.24E-02 |
| 3a_hand - V4t | -0.21 | 0.04 | 756 | -5.86 | 1.38E-05 |
| 3a_hand - V7 | -0.19 | 0.04 | 756 | -5.35 | 2.31E-04 |
| 3a_hand - VVC | -0.17 | 0.04 | 756 | -4.87 | 2.78E-03 |
| 3b_hand - 55b | -0.17 | 0.04 | 756 | -4.92 | 2.18E-03 |
| 3b_hand - 6v | -0.23 | 0.04 | 756 | -6.49 | 3.16E-07 |
| 3b_hand - a24pr | -0.21 | 0.04 | 756 | -6.03 | 5.09E-06 |
| 3b_hand - FFC | -0.19 | 0.04 | 756 | -5.30 | 3.07E-04 |
| 3b_hand - LO2 | -0.22 | 0.04 | 756 | -6.38 | 6.33E-07 |
| 3b_hand - MT | -0.18 | 0.04 | 756 | -5.01 | 1.38E-03 |
| 3b_hand - OP1 | -0.18 | 0.04 | 756 | -5.05 | 1.14E-03 |
| 3b_hand - p24pr | -0.20 | 0.04 | 756 | -5.78 | 2.15E-05 |
| 3b_hand - PGp | -0.16 | 0.04 | 756 | -4.56 | 1.18E-02 |
| 3b_hand - PIT | -0.23 | 0.04 | 756 | -6.55 | 2.19E-07 |
| 3b_hand - SFL | -0.18 | 0.04 | 756 | -4.99 | 1.52E-03 |
| 3b_hand - TE2p | -0.16 | 0.04 | 756 | -4.65 | 7.80E-03 |
| 3b_hand - TPOJ2 | -0.17 | 0.04 | 756 | -4.72 | 5.69E-03 |
| 3b_hand - TPOJ3 | -0.16 | 0.04 | 756 | -4.47 | 1.84E-02 |
| 3b_hand - V3CD | -0.18 | 0.04 | 756 | -5.19 | 5.41E-04 |
| 3b_hand - V4t | -0.23 | 0.04 | 756 | -6.63 | 1.30E-07 |
| 3b_hand - V7 | -0.21 | 0.04 | 756 | -6.12 | 3.01E-06 |
| 3b_hand - V8 | -0.15 | 0.04 | 756 | -4.30 | 3.90E-02 |
| 3b_hand - VVC | -0.20 | 0.04 | 756 | -5.63 | 5.00E-05 |
| 4_face - 6v | -0.16 | 0.04 | 756 | -4.43 | 2.18E-02 |
| 4_face - LO2 | -0.15 | 0.04 | 756 | -4.32 | 3.59E-02 |
| 4_face - PIT | -0.16 | 0.04 | 756 | -4.49 | 1.67E-02 |
| 4_face - V4t | -0.16 | 0.04 | 756 | -4.57 | 1.15E-02 |
| 6a - 6v | -0.17 | 0.04 | 756 | -4.75 | 5.02E-03 |
| 6a - a24pr | -0.15 | 0.04 | 756 | -4.29 | 4.09E-02 |
| 6a - LO2 | -0.16 | 0.04 | 756 | -4.63 | 8.53E-03 |
| 6a - PIT | -0.17 | 0.04 | 756 | -4.80 | 3.80E-03 |
| 6a - V4t | -0.17 | 0.04 | 756 | -4.88 | 2.55E-03 |
| 6a - V7 | -0.15 | 0.04 | 756 | -4.38 | 2.76E-02 |
| 6mp - 6v | -0.17 | 0.04 | 756 | -4.92 | 2.14E-03 |

| | | | | | |
|---|---|---|---|---|---|
| 6mp - a24pr | -0.16 | 0.04 | 756 | -4.46 | 1.87E-02 |
| 6mp - LO2 | -0.17 | 0.04 | 756 | -4.81 | 3.70E-03 |
| 6mp - PIT | -0.17 | 0.04 | 756 | -4.98 | 1.60E-03 |
| 6mp - V4t | -0.18 | 0.04 | 756 | -5.06 | 1.06E-03 |
| 6mp - V7 | -0.16 | 0.04 | 756 | -4.55 | 1.25E-02 |
| 6v - 8BM | 0.19 | 0.04 | 756 | 5.39 | 1.87E-04 |
| 6v - IP0 | 0.17 | 0.04 | 756 | 4.74 | 5.05E-03 |
| 6v - IP2 | 0.18 | 0.04 | 756 | 5.10 | 8.71E-04 |
| 6v - MIP | 0.16 | 0.04 | 756 | 4.57 | 1.16E-02 |
| 6v - PFt | 0.17 | 0.04 | 756 | 4.95 | 1.86E-03 |
| 8BM - a24pr | -0.17 | 0.04 | 756 | -4.94 | 1.97E-03 |
| 8BM - LO2 | -0.19 | 0.04 | 756 | -5.28 | 3.39E-04 |
| 8BM - p24pr | -0.16 | 0.04 | 756 | -4.69 | 6.60E-03 |
| 8BM - PIT | -0.19 | 0.04 | 756 | -5.45 | 1.37E-04 |
| 8BM - V4t | -0.19 | 0.04 | 756 | -5.53 | 8.79E-05 |
| 8BM - V7 | -0.18 | 0.04 | 756 | -5.02 | 1.27E-03 |
| 8BM - VVC | -0.16 | 0.04 | 756 | -4.54 | 1.33E-02 |
| a24pr - IP0 | 0.15 | 0.04 | 756 | 4.29 | 4.11E-02 |
| a24pr - IP2 | 0.16 | 0.04 | 756 | 4.64 | 8.17E-03 |
| a24pr - PFt | 0.16 | 0.04 | 756 | 4.49 | 1.64E-02 |
| IP0 - LO2 | -0.16 | 0.04 | 756 | -4.63 | 8.59E-03 |
| IP0 - PIT | -0.17 | 0.04 | 756 | -4.80 | 3.82E-03 |
| IP0 - V4t | -0.17 | 0.04 | 756 | -4.88 | 2.57E-03 |
| IP0 - V7 | -0.15 | 0.04 | 756 | -4.38 | 2.78E-02 |
| IP2 - LO2 | -0.18 | 0.04 | 756 | -4.99 | 1.53E-03 |
| IP2 - p24pr | -0.15 | 0.04 | 756 | -4.39 | 2.56E-02 |
| IP2 - PIT | -0.18 | 0.04 | 756 | -5.16 | 6.47E-04 |
| IP2 - V4t | -0.18 | 0.04 | 756 | -5.24 | 4.24E-04 |
| IP2 - V7 | -0.17 | 0.04 | 756 | -4.73 | 5.37E-03 |
| IP2 - VVC | -0.15 | 0.04 | 756 | -4.24 | 4.96E-02 |
| LO2 - MIP | 0.16 | 0.04 | 756 | 4.46 | 1.93E-02 |
| LO2 - PFt | 0.17 | 0.04 | 756 | 4.84 | 3.23E-03 |
| MIP - PIT | -0.16 | 0.04 | 756 | -4.63 | 8.81E-03 |
| MIP - V4t | -0.17 | 0.04 | 756 | -4.71 | 5.99E-03 |
| p24pr - PFt | 0.15 | 0.04 | 756 | 4.24 | 4.99E-02 |
| PFt - PIT | -0.18 | 0.04 | 756 | -5.01 | 1.39E-03 |
| PFt - V4t | -0.18 | 0.04 | 756 | -5.09 | 9.23E-04 |
| PFt - V7 | -0.16 | 0.04 | 756 | -4.58 | 1.09E-02 |

Supplementary table 27: **Nuggets across ROIs.** Two-sided t contrasts between ROIs (Bonferroni-corrected) with Kenward-Roger's method for degrees of freedom. Contrast estimates are expressed as difference of square-root scaled values. Only significant contrasts are reported.

| contrast | estimate | SE | df | t.ratio | p.value |
|---|---|---|---|---|---|
| V4 - AVI | -0.71 | 0.15 | 756 | -4.64 | 8.25E-03 |
| V4 - IFJp | -0.72 | 0.15 | 756 | -4.72 | 5.55E-03 |
| V4 - OP1 | -0.68 | 0.15 | 756 | -4.44 | 2.09E-02 |
| V4 - p9_46v | -0.66 | 0.15 | 756 | -4.30 | 3.90E-02 |
| V4 - PEF | -0.72 | 0.15 | 756 | -4.73 | 5.38E-03 |
| V4 - V7 | -0.68 | 0.15 | 756 | -4.44 | 2.07E-02 |
| 3a_hand - 4_face | 0.67 | 0.15 | 756 | 4.37 | 2.90E-02 |
| 3b_hand - IFJp | -0.66 | 0.15 | 756 | -4.29 | 4.12E-02 |
| 3b_hand - PEF | -0.66 | 0.15 | 756 | -4.29 | 4.00E-02 |
| 4_face - AVI | -0.79 | 0.15 | 756 | -5.16 | 6.21E-04 |
| 4_face - FEF | -0.66 | 0.15 | 756 | -4.33 | 3.48E-02 |
| 4_face - IFJp | -0.80 | 0.15 | 756 | -5.25 | 4.02E-04 |
| 4_face - IPS1 | -0.65 | 0.15 | 756 | -4.26 | 4.69E-02 |
| 4_face - MIP | -0.72 | 0.15 | 756 | -4.71 | 5.89E-03 |
| 4_face - MT | -0.70 | 0.15 | 756 | -4.56 | 1.18E-02 |
| 4_face - OP1 | -0.76 | 0.15 | 756 | -4.96 | 1.72E-03 |
| 4_face - p9_46v | -0.74 | 0.15 | 756 | -4.82 | 3.44E-03 |
| 4_face - PEF | -0.80 | 0.15 | 756 | -5.26 | 3.88E-04 |
| 4_face - TPOJ2 | -0.65 | 0.15 | 756 | -4.26 | 4.74E-02 |
| 4_face - V7 | -0.76 | 0.15 | 756 | -4.97 | 1.71E-03 |
| AVI - FFC | 0.65 | 0.15 | 756 | 4.25 | 4.76E-02 |
| AVI - TE2p | 0.65 | 0.15 | 756 | 4.27 | 4.44E-02 |
| AVI - VVC | 0.67 | 0.15 | 756 | 4.37 | 2.80E-02 |
| FFC - IFJp | -0.66 | 0.15 | 756 | -4.34 | 3.30E-02 |
| FFC - PEF | -0.67 | 0.15 | 756 | -4.34 | 3.20E-02 |
| IFJp - TE2p | 0.67 | 0.15 | 756 | 4.35 | 3.07E-02 |
| IFJp - VVC | 0.68 | 0.15 | 756 | 4.46 | 1.92E-02 |
| PEF - TE2p | 0.67 | 0.15 | 756 | 4.36 | 2.98E-02 |
| PEF - VVC | 0.68 | 0.15 | 756 | 4.46 | 1.86E-02 |

Supplementary table 28: **Ranges across ROIs.** Two-sided t contrasts between ROIs (Bonferroni-corrected) with Kenward-Roger's method for degrees of freedom. Contrast estimates are expressed as difference of log scaled values. Only significant contrasts are reported.

### Supplementary Methods - MRI data processing

Results included in this manuscript come from preprocessing performed using *fMRIPrep* 21.0.2 ([1]; [2]; RRID:SCR_016216), which is based on *Nipype* 1.6.1 ([3]; [4]; RRID:SCR_002502).

#### Preprocessing of B0 inhomogeneity mappings

A total of 12 fieldmaps were found available within the input BIDS structure for this particular subject. A *B0*-nonuniformity map (or *fieldmap*) was estimated based on two (or more) echo-planar imaging (EPI) references with `topup` ([5]; FSL 6.0.5.1:57b01774).

#### Anatomical data preprocessing

A total of 1 T1-weighted (T1w) images were found within the input BIDS dataset. The T1-weighted (T1w) image was corrected for intensity non-uniformity (INU) with `N4BiasFieldCorrection` [6], distributed with ANTs 2.3.3 [7, RRID:SCR_004757], and used as T1w-reference throughout the workflow. The T1w-reference was then skull-stripped with a *Nipype* implementation of the `antsBrainExtraction.sh` workflow (from ANTs), using OASIS30ANTs as target template. Brain tissue segmentation of cerebrospinal fluid (CSF), white-matter (WM) and gray-matter (GM) was performed on the brain-extracted T1w using `fast` [FSL 6.0.5.1:57b01774, RRID:SCR_002823, 8]. Brain surfaces were reconstructed using `recon-all` [FreeSurfer 6.0.1, RRID:SCR_001847, 9], and the brain mask estimated previously was refined with a custom variation of the method to reconcile ANTs-derived and FreeSurfer-derived segmentations of the cortical gray-matter of Mindboggle [RRID:SCR_002438, 10]. Volume-based spatial normalization to two standard spaces (MNI152NLin6Asym, MNI152NLin2009cAsym) was performed through nonlinear registration with `antsRegistration` (ANTs 2.3.3), using brain-extracted versions of both T1w reference and the T1w template. The following templates were selected for spatial normalization: *FSL's MNI ICBM 152 non-linear 6th Generation Asymmetric Average Brain Stereotaxic Registration Model*[[11], RRID:SCR_002823; TemplateFlow ID: MNI152NLin6Asym], *ICBM 152 Nonlinear Asymmetrical template version 2009c*[[12], RRID:SCR_008796; TemplateFlow ID: MNI152NLin2009cAsym].

#### Functional data preprocessing

For each of the 12 BOLD runs found per subject (across all tasks and sessions), the following preprocessing was performed. First, a reference volume and its skull-stripped version were generated using a custom methodology of *fMRIPrep*. Head-motion parameters with respect to the BOLD reference (transformation matrices, and six corresponding rotation and translation parameters) are estimated before any spatiotemporal filtering using `mcflirt` [FSL 6.0.5.1:57b01774, 13]. The estimated *fieldmap* was then aligned with rigid-registration to the target EPI (echo-planar imaging) reference run. The field coefficients were mapped on to the reference EPI using the transform. The BOLD reference was then co-registered to the T1w reference using `bbregister` (FreeSurfer) which implements boundary-based registration [14]. Co-registration was configured with six degrees of freedom. Several confounding time-series were calculated based on the *preprocessed BOLD*: framewise displacement (FD), DVARS and three region-wise global signals. FD was computed using two formulations following Power (absolute sum of relative motions, [15]) and Jenkinson (relative root mean square displacement between affines, [13]). FD and DVARS are calculated for each functional run, both using their implementations in *Nipype* [following the definitions by 15]. The three global signals are extracted within the CSF, the WM, and the whole-brain masks. Additionally, a set of physiological regressors were extracted to allow for component-based noise correction [*CompCor*, 16]. Principal components are estimated after high-pass filtering the *preprocessed BOLD* time-series (using a discrete cosine filter with 128s cut-off) for the two *CompCor* variants: temporal (tCompCor) and anatomical (aCompCor). tCompCor components are then calculated from the top 2% variable voxels within the brain mask. For aCompCor, three probabilistic masks (CSF, WM and combined CSF+WM) are generated in anatomical space. The implementation differs from that of Behzadi et al. in that instead of eroding the masks by 2 pixels on BOLD space, the aCompCor masks are subtracted a mask of pixels that likely contain a volume fraction of GM. This mask is obtained by dilating a GM mask extracted from the FreeSurfer's *aseg* segmentation, and it ensures components are not extracted from voxels containing a minimal fraction of GM. Finally, these masks are resampled into BOLD space and binarized by thresholding at 0.99 (as in the original implementation). Components are also calculated separately within the WM and CSF masks. For each CompCor decomposition, the $k$ components with the largest singular values are retained, such that the retained components' time series are sufficient to explain 50 percent of variance across the nuisance mask (CSF, WM, combined, or temporal). The remaining components are dropped from consideration. The head-motion estimates calculated in the correction step were also placed within the corresponding confounds file. The confound time series derived from head motion estimates and global signals were expanded with the inclusion of temporal derivatives and quadratic terms for each [17]. Frames that exceeded a threshold of 0.5 mm FD or 1.5 standardised DVARS were annotated as motion outliers. The BOLD time-series were resampled into standard space, generating a *preprocessed BOLD run in MNI152NLin6Asym space*. First, a reference volume and its skull-stripped version were generated using a custom methodology of *fMRIPrep*. The BOLD time-series were resampled onto the following surfaces (FreeSurfer reconstruction nomenclature): *fsnative*, *fsaverage*. Automatic removal of motion artifacts using independent component analysis [ICA-AROMA, 18] was performed on the *preprocessed BOLD on MNI space* time-series after removal of non-steady state volumes and spatial smoothing with an isotropic, Gaussian kernel of 6mm FWHM (full-width half-maximum). Corresponding "non-aggresively" denoised runs were produced after such smoothing. Additionally, the "aggressive" noise-regressors were collected and placed in the corresponding confounds file. All resamplings can be performed with *a single interpolation step* by composing all the pertinent transformations (i.e. head-motion transform matrices, susceptibility distortion correction when available, and co-registrations to anatomical and output spaces). Gridded (volumetric) resamplings were performed using `antsApplyTransforms` (ANTs), configured with Lanczos interpolation to minimize the smoothing effects of other kernels [19]. Non-gridded (surface) resamplings were performed using `mri_vol2surf` (FreeSurfer).

Many internal operations of *fMRIPrep* use *Nilearn* 0.8.1 [20, RRID:SCR_001362], mostly within the functional processing workflow. For more details of the pipeline, see the section corresponding to workflows in *fMRIPrep*'s documentation.

##### Copyright Waiver

The above boilerplate text was automatically generated by fMRIPrep with the express intention that users should copy and paste this text into their manuscripts *unchanged*. It is released under the CC0 license.

[13] M. Jenkinson, P. Bannister, M. Brady, and S. Smith, "Improved optimization for the robust and accurate linear registration and motion correction of brain images," *NeuroImage*, vol. 17, no. 2, pp. 825–

841, 2002, ISSN: 1053-8119. DOI: 10.1006/nimg.2002.1132. [Online]. Available: http://www.sciencedirect.com/science/article/pii/S1053811902911328.
